# Electrostatic interactions in nucleosome and higher-order structures are regulated by protonation state of histone ionizable residue

**DOI:** 10.1101/2024.06.07.597724

**Authors:** Houfang Zhang, Wenhan Guo, Wang Xu, Anbang Li, Lijun Jiang, Lin Li, Yunhui Peng

**Affiliations:** Institute of Biophysics and Department of Physics, Central China Normal University, Wuhan, 430079, China; Department of Physics, University of Texas at El Paso, El Paso, TX 79902, USA; Hubei Key Laboratory of Genetic Regulation & Integrative Biology, School of Life Sciences, Central China Normal University, Wuhan 430079, China

**Keywords:** chromatin structure, nucleosome, pKa shift, proton transfer, electrostatic interaction, histone cancer mutation

## Abstract

The nucleosome serves as the fundamental unit of chromatin organization, with electrostatic interactions acting as the driving forces in the folding of nucleosomes into chromatin. Perturbations in cellular pH conditions can lead to changes in the protonation states of titratable histone residues, impacting nucleosome surface electrostatic potentials and interactions. However, the effects of proton uptake or release of histone ionizable groups on nucleosome-partner protein interactions and higher-order chromatin structures remain largely unexplored. Here, we conducted comprehensive analyses of histone titratable residue pKa values in various nucleosome contexts, utilizing 96 experimentally determined structures. We revealed that pH-induced changes in histone residue protonation states modulated nucleosome surface electrostatic potentials and significantly influenced nucleosome-partner protein interactions. Furthermore, we observed that proton uptake or release often accompanied nucleosome-partner protein interactions, facilitating their binding processes. Additionally, using a dataset of 1266 recurrent histone cancer mutations, we systematically characterized their impact on nucleosome surface electrostatics, demonstrating their profound effects on electrostatic interactions between nucleosomes and partner proteins. Finally, our findings suggest that alterations in histone protonation or cancer mutations can also regulate nucleosome self-association, thereby modulating the organization and dynamics of higher-order chromatin structure.

## Introduction

In eukaryotic cells, the nucleosome serves as the fundamental unit of chromatin, playing a crucial role in packaging and organizing DNA within the nucleus^1, 2^. The nucleosome core particle (NCP) consists of approximately 146-148 base pairs of DNA molecules wrapped around a histone octamer, which comprises two copies of each of the four histone proteins: H2A, H2B, H3, and H4^3, 4^. The compaction of nucleosomes into higher-order structures involves the stacking of nucleosomes on top of each other, facilitated by the flexible linker DNA between nucleosomes and the bridging of tails between adjacent NCPs^5–7^. Given the polyelectrolyte nature of nucleosome and linker DNA, electrostatic interactions are inherently the driving forces in the folding of arrays of nucleosomes into chromatin^8, 9^.

DNA is one of the most highly charged polymers in cells, generating a strong negative electrostatic field. To compact DNA, the electrostatic self-repulsion between DNA molecules needs to be overcome^10, 11^. Thus, the association of the DNA around the positively-charged histone octamer would help the electrostatic screening of negative charges on the DNA backbones. It has long been assumed that formation of nucleosome structure results in completely electrostatic screening of the DNA and a weak electrostatic field surrounding the nucleosome^12^. However, recent studies have experimentally shown that despite the attenuation of charges in the complexation of the DNA into the nucleosome, nucleosome still maintains a strong, negative electrostatic field and it is important to consider the essential roles of polyelectrolyte nature of the NCPs in nucleosome compaction and interaction^12, 13^.

Experimental evidence has demonstrated that the protonation states of titratable histone residues near the acidic patch can change within the physiological pH range, thereby affecting the nucleosome surface electrostatic potential^14^. Biologically relevant fluctuations in intracellular pH can significantly impact the stability of nucleosome structures by altering the protonation states of titratable groups^15^. However, it is still not well understood whether and how such changes in nucleosome electrostatics could modulate the nucleosome-nucleosome interactions and higher-order nucleosome structures.

The formation of protein complexes often entails changes in pKa values and protonation states of ionizable groups, driven by direct electrostatic interactions, structural rearrangements of receptors, and desolvation of the groups involved protein binding^16–18^. Nucleosomes interact with a plethora of chromatin factors, including chromatin-modifying enzymes, chromatin remodelers, and transcription factors^19–21^. Given the polyelectrolyte nature of nucleosomes, it is reasonable to expect that the binding of proteins to nucleosomes frequently involves the proton uptake or release of ionizable groups. However, the underlying mechanisms of proton uptake or release in histone interactions and their effects on nucleosome-chromatin factor interactions were largely understudied.

Numerous recent studies have underscored the essential roles of histone mutations in the oncogenic development of various cancer types^22, 23^ and histone cancer mutations have profound effects on nucleosome dynamics, interactions, and higher-order chromatin structures ^24, 25^. Our recent computational analysis has further indicated that histone cancer mutations are enriched on histone binding interfaces and have strongly disruptive effects on histone-histone, histone-DNA, and histone-partner protein interactions^26^. These mutations can dramatically alter the physicochemical properties of the histone binding interface, suppressing both positive and negative net charges on the surface of the histone octamer^26^. These alterations may further affect nucleosome surface electrostatics and long-range electrostatic interactions, potentially leading to profound effects on nucleosome-partner protein interactions and higher-order chromatin structures.

Here, we conducted a systematic analysis of the pKa values of histone titratable residues in free dimers, nucleosomes, and nucleosome complex structures using a diverse array of experimentally determined structures. Through structural analyses and electrostatic calculations, we discovered that alterations in protonation states of key histone residues can occur due to perturbations in pH near cellular physiological conditions, resulting in significant effects on nucleosome surface electrostatic potentials and nucleosome-partner protein interactions. Additionally, through molecular dynamics simulations and free energy calculations, we demonstrated that the binding of regulatory proteins to nucleosomes was coupled with proton uptake or release of histone titratable residues, generally promoting nucleosome-partner protein interactions. Furthermore, we systematically characterized the impact of histone cancer mutations on nucleosome surface electrostatics, revealing profound effects on the strengths and directions of electrostatic forces between nucleosome and partner proteins. Finally, we explored the effects of changes in histone residue protonation and cancer mutations on nucleosome stacking and H4 tail-acidic patch interactions. Our results suggest that alterations in histone protonation or cancer mutations may not only impact individual nucleosomes but also influence nucleosome self-association and higher-order nucleosome structure, thereby contributing to the overall organization and dynamics of chromatin structure.

## Methods

### Collection of nucleosome complex structures

To analyze the electrostatic interactions between nucleosome and various regulatory proteins, we collected all available nucleosome complex structures from RCSB PDB bank^27^, a total of 234 nucleosome structures up to the date of this paper submitted. Utilizing our developed histone interaction network, which encompasses 3808 histone-regulatory protein interactions^24, 28^, we systematically characterized various types of histone binding interfaces. The interfacial residues were determined by residues located within a 5 Å distance between heavy atoms of histone and their respective binding partner. To eliminate redundancy, where the same nucleosome binding proteins may have multiple available complex structures, we selected representative structures for each binding protein following the criteria outlined in our recent study^24, 28^. Ultimately, 83 representative nucleosome complex structures were chosen for subsequent pKa calculations and electrostatic potential analyses (Supplementary Table 1 and Supplementary Figure 1).

### pKa calculations of nucleosome structures

To evaluate the influence of protein structure environment on histone residue protonation states, we computed the pKa values of histone titratable residues within histone dimers (H2A-H2B and H3-H4) and within the context of nucleosome structures. We selected 13 high-resolution nucleosome structures (Resolution < 3 Å) for analysis, from three different organisms: Homo sapiens, Xenopus laevis, and Drosophila melanogaster (Supplementary Table 2). In each selected nucleosome structure, we extracted each copy of the H2A-H2B and H3-H4 histone dimers and subjected them to 20,000 steps of energetic minimization using Amber ^29^ with the ff14SB force field for proteins and OL15 for DNA. Subsequently, the minimized dimer structures were utilized for pKa calculations of histone titratable residues (Glu, Asp, Lys, His, and Arg) using PROPKA^30^ (Figure 1a). To analyze the pKa changes attributable to the protein environment in nucleosomes, the pKa values of histone titratable residues were computed using the corresponding nucleosome structures. The pKa shift of each histone residue was determined as follows:

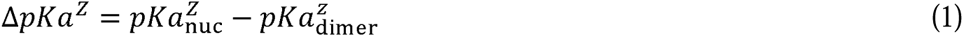

, where Z denotes the titratable group (Glu, Asp, Lys, His, and Arg) in histones, and ΔpKa represents the difference in pKa values of histone residues between the dimer and the octamer within the nucleosome structure.

**Figure 1.**
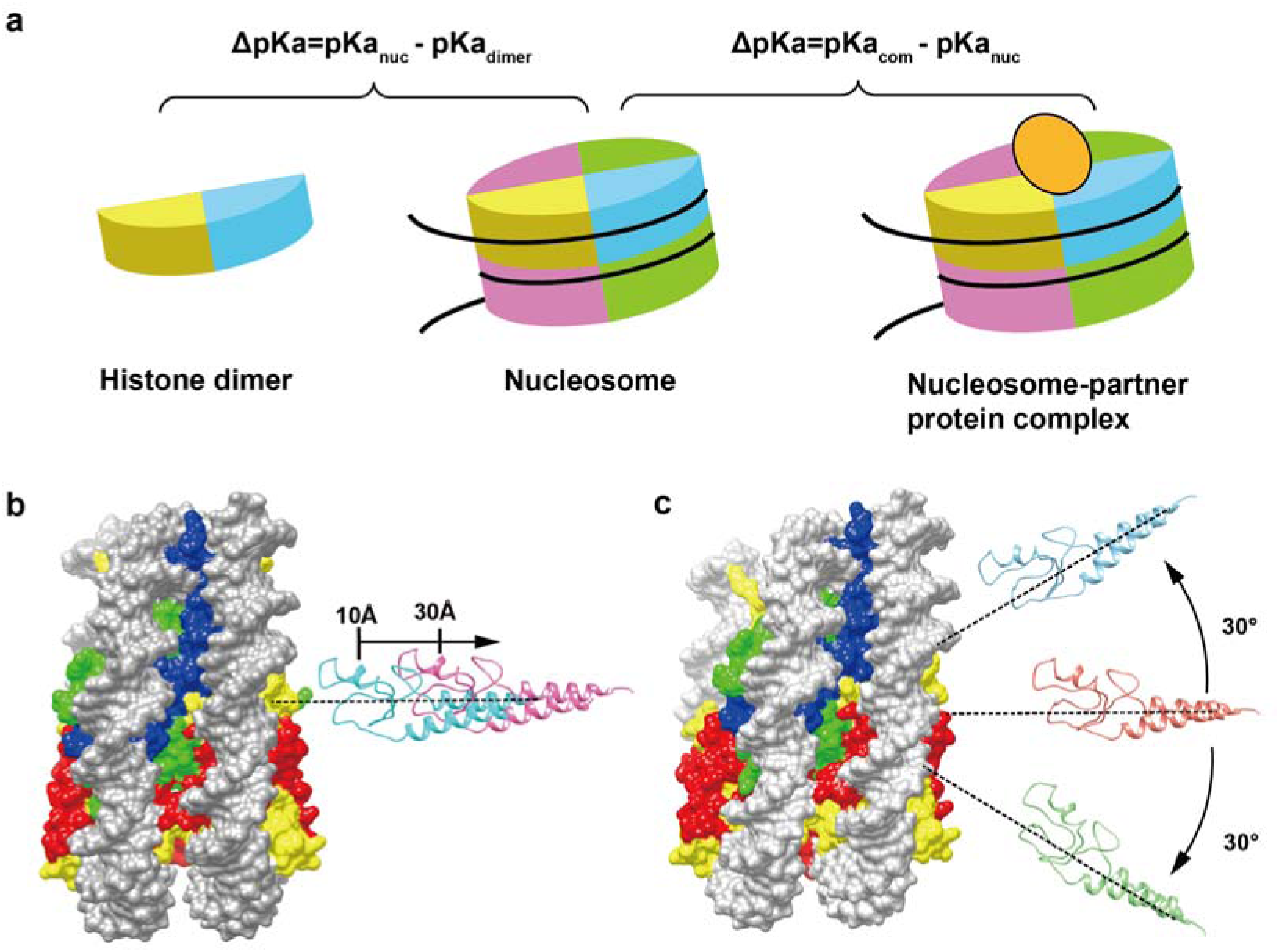
Schematic diagram of pKa calculations and electrostatic analyses of nucleosome structures. (a) Determination of pKa shifts in histone titratable residues induced by the protein environment in nucleosome or the nucleosome-partner protein interactions. (b) Calculation of the long-range electrostatic forces generated between nucleosomes and binding proteins at separating distances from 10 to 30 Å, with a step size of 2 Å, along the line connecting their centers of mass. (c) Calculation of the long-range electrostatic forces generated between nucleosomes and binding proteins at different rotational angles of the line connecting their centers of mass, from −30° to 30°, with a step size of 6°.

It is well-known that protein-protein and protein-nucleic interactions are associated with pKa shifts and changes in the protonation states of titratable residues^18, 31^. Here, we systematically analyzed pKa shifts and protonation state changes of histone residues induced by the interaction between regulatory proteins and nucleosomes. We performed pKa calculations of histone titratable groups for 83 representative nucleosome complex structures using PROPKA3^30^ (Figure 1a). For each nucleosome complex structure, we initially computed the pKa values of protein residues within the nucleosome complex (bound state). Subsequently, we calculated the pKa values with the individual nucleosome structure (unbound state). The pKa shift (ΔpKa) for each titratable group induced by the interaction between nucleosome and binding protein was then determined using the following equation:

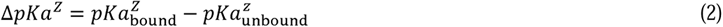

where Z represents the titratable group (Glu, Asp, Lys, His, and Arg) in histones or nucleosome-binding proteins, and ΔpKa represents the change in pKa value of the titratable group from the unbound state to the bound state.

To analyze the pH-induced changes in protonation states of histone titratable residues, we categorized these residues into four classes based on the pKa calculations: i) Titratable groups experiencing changes in their protonation states within the physiological pH range (pH 6.5 to 7.5). ii) Titratable groups experiencing changes in protonation states under slightly acidic condition (pH 5 to 6.5). iii) Titratable groups experiencing changes in protonation states under weakly basic condition (pH 7.5 to 9). iv) Titratable groups that do not experience changes in their protonation states within the physiological or mildly acidic/basic pH ranges (5 < pH < 9, less sensitive to changes in pH within the typical cellular environment).

### Electrostatic potential analyses of nucleosome structures

The electrostatic potentials of nucleosome structures were calculated using the Delphi program^32, 33^, which solves the Poisson-Boltzmann equation (PBE) with the finite difference method:

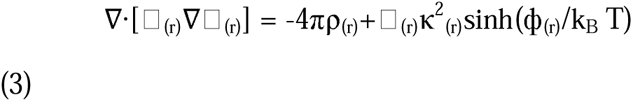

ρ(r) represents the permanent charge density, ε(r) denotes the dielectric permittivity, T signifies temperature, κ stands for the Debye-Huckel parameter, and k_B_ represents the Boltzmann constant.

In our calculations, we set the dielectric constants for protein and solvent to 2 and 80, respectively, and the salt concentration constant was maintained at 0.15 M. The percentage filling of the box was adjusted to 70 using a scale of 1 grid/Å and a water probe radius of 1.4 Å. We utilized amber parameter files for atomic charge and radius information. The resulting electrostatic potential map was saved in CUBE format and subsequently visualized and analyzed using UCSF Chimera^34^.

### Electrostatic force calculations between nucleosome and binding proteins

The long-range electrostatic forces between nucleosomes and binding proteins were computed using DelphiForce^35^, a tool that derives the magnitudes and directions of electrostatic forces by solving the negative gradient of electrostatic potentials. Utilizing the finite difference method, the electrostatic potential Φ(r) is obtained at each grid point (i, j, k), denoted as Φ(i, j, k), and the components of the electrostatic forces (E(i, j, k)) are calculated as follows:

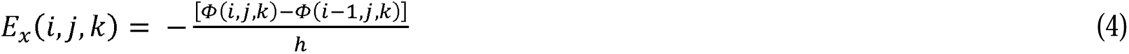

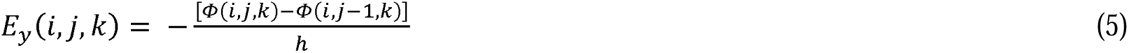

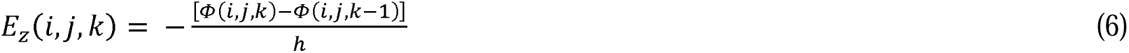

, where *i*, *j*, and *k* are the indices of finite difference grids in the *x*, *y*, *z* directions, and *h* is the distance between two neighboring grids.

To calculate the long-range electrostatic forces generated between nucleosome and binding protein, the binding protein in each nucleosome complex structure were separated away at certain distances from histone octamer in the direction of the line that connects their centers of mass using StructureMan^36^ (Figure 1). Subsequently, the electrostatic forces between the nucleosome and binding proteins were computed at different separating distances of their centers of mass from 10 to 30 Å with a step size of 2 Å, and at different rotation angles of the line connecting their centers of mass from −30° to 30° with a step size of 6°.

### Molecular dynamics simulations of nucleosome complex structures under different protonation states of histone ionizable groups

Given that the binding of proteins to nucleosomes induces changes in the protonation states of key histone titratable residues, we conducted molecular dynamics simulations of nucleosome complex structures to explore how these changes could influence nucleosome-partner interactions. For our simulations, we chose seven representative nucleosome complex structures containing various nucleosome-binding proteins involved in epigenetic regulation processes, including regulatory protein SIR3, tumor suppressor p53-binding protein 1, SWI/SNF chromatin remodeler RSC, and DNA methyltransferase.

For each nucleosome complex structure, we conducted two sets of simulations (Supplementary Table 3). In one set, histone residues were assigned protonation states corresponding to the unbound condition, while in the other set, histone residues were assigned protonation states reflecting the bound condition. By comparing the results of these simulations, our objective is to evaluate the influence of changes in the protonation states of histone residues on nucleosome-partner protein interactions.

All simulations were conducted under the AMBER force field (ff14SB force field for proteins and OL15 for DNA molecules) and employing the TIP4PEW water model in explicit solvent. In each simulation system, the initial structural model was solvated with 0.15 M NaCl in a cubic water box, maintaining a minimum distance of 10 Å from the atoms of the nucleosome complex to the edge of water box. The systems were kept at T=310 K using Langevin dynamics with a constant pressure simulation at 1 atm. SHAKE bond length constraints were applied for bonds involving hydrogens, with a non-bonded interaction cutoff distance set at 10 Å. Electrostatic calculations were performed using the Particle Mesh Ewald (PME) method, employing a spacing of 1 Å and a real-space cutoff of 12 Å. Periodic boundary conditions were applied, and trajectories were saved every 10 ps. Each system underwent 10,000 steepest descent minimizations followed by another 10,000 conjugate gradient minimizations. After minimization, the systems were gradually heated from 100 to 310 K in the NVT ensemble, followed by switching to the NPT ensemble for 2ns equilibrations before 100 ns production runs.

The binding free energies between the nucleosome and binding partners were calculated using the molecular mechanics Poisson-Boltzmann surface area (MM/PBSA) method implemented in the Amber22 package^29^. Calculations were performed for every 100 ps frame, excluding the first 20 ns trajectories. Residue-wise decomposition was then applied to derive the binding energy per histone residue. The hydrogen bond and salt bridge in histone binding interfaces were analyzed using the customized VMD TCL scripts.

### Analysis of histone cancer mutations

To examine the impact of histone cancer mutations on nucleosome electrostatic potentials and long-range electrostatic interactions, we utilized one recently compiled dataset of histone cancer-associated mutations, which encompasses 7940 mutations across 84 histone genes and 83 cancer types^24^. Subsequently, we filtered out mutations previously documented in the dbSNP database^37^ and the mutations from samples with a tumor mutation burden (TMB) of > 10 mutations per megabase (Mb) following previous study^38^ and identified recurrent histone mutations, indicative of potential associations with cancer progression. Recurrent histone cancer mutations were defined as those occurring in at least two distinct patients within the dataset. Following this filtering process, we curated a refined dataset, termed the ‘refined histone cancer mutation set’, which comprises 1266 recurrent missense mutations across 77 histone genes and 55 cancer types. This refined dataset was then subjected to further electrostatic analyses in this study.

To assess the impact of mutation on nucleosome-partner protein interactions, we mapped histone cancer mutations onto our recently developed histone-partner protein interaction networks^26^. Then, the changes in protein binding free energy (ΔΔGs) caused by each mutation were calculated using the SAAMBE-3D approach^39^, a newly developed machine learning algorithm to predict the effects of single amino acid substitutions on protein-protein interactions (PPIs).

## Results

### Histone ionizable residues in nucleosome undergo substantial pKa changes due to protein environments

Wrapping of DNA molecules around histone octamers creates a unique protein environment in nucleosome structure, which can significantly influence the pKa values of histone residues and further impact the ionizable states of histone residues even at physiological pH condition. To systematically characterize the pKa changes and corresponding ionizable states of histone residues, we have performed structured-based pKa calculations of all histone titratable residues using 13 high-resolution nucleosome structures from three organisms (Supplementary Table 2). Moreover, the pKa values were further predicted using the corresponding refined H2A-H2B and H3-H4 dimer structures and the pKa changes of each histone titratable residues caused by the protein environment in nucleosome were derived using equation 1 (see methods section and Figure 1). To validate our predictions, we have further compared our predicted pKa values with experimental determined pKa values of nine titratable residues near the acidic patch^40^, which achieved a Pearson correlation coefficient of 0.85 (Figure 2a and Supplementary Table 4).

**Figure 2.**
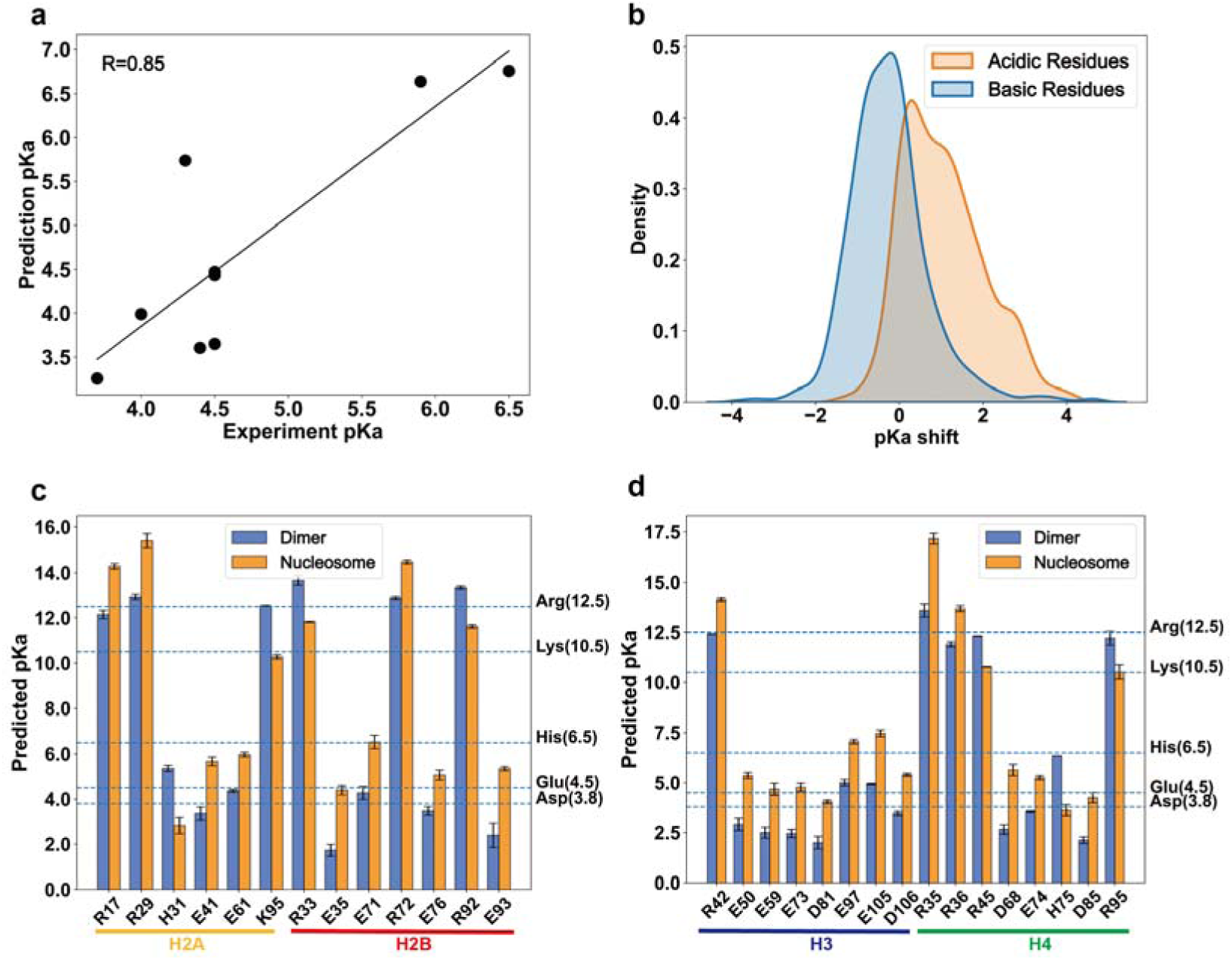
pKa shifts of histone titratable residues from histone dimer to octamer formation. (a) Comparison of our predicted pKa values with experimentally determined pKa values for nine titratable residues near the acidic patch. (b) Distribution of pKa shifts of histone titratable residues due to the protein environments in nucleosome. The pKa shift represents the difference in pKa values of histone titratable residues between the dimer and the octamer within the nucleosome structure. (c) and (d) Histone titratable residues exhibiting significant pKa shifts in nucleosome structures (|ΔpKa| ≥ 1.5). Histone residues belonging to different histone types were indicated by different colors.

Through our analysis of pKa changes, we observed that more than 60% of histone basic residues (Arg, Lys, and His) displayed decreased pKa values in nucleosomes compared to free dimers (Figure 2b). In contrast, over 90% of histone acidic groups (Asp and Glu) showed elevated pKa values, indicating suppressed acidity. This suggests a generally weakened basicity and acidity of histone residues within the nucleosome structure.

In addition, we identified 16 acidic residues and 13 basic residues exhibiting significant pKa shifts in at least three different nucleosome structures (|ΔpKa| ≥ 1.5), with the largest changes up to 4 (Figure 2). All of these residues were located on the histone-histone and/or histone-DNA binding interfaces, suggesting substantial changes in their local environment from histone dimer to octamer formation (Supplementary Tables 5 and 6). Histone residues with significant pKa shifts were distributed among all four histone types and were predominantly situated around the acidic patch, H3 α1L1 elbow, and H3 alpha2 (Figure 3a).

**Figure 3.**
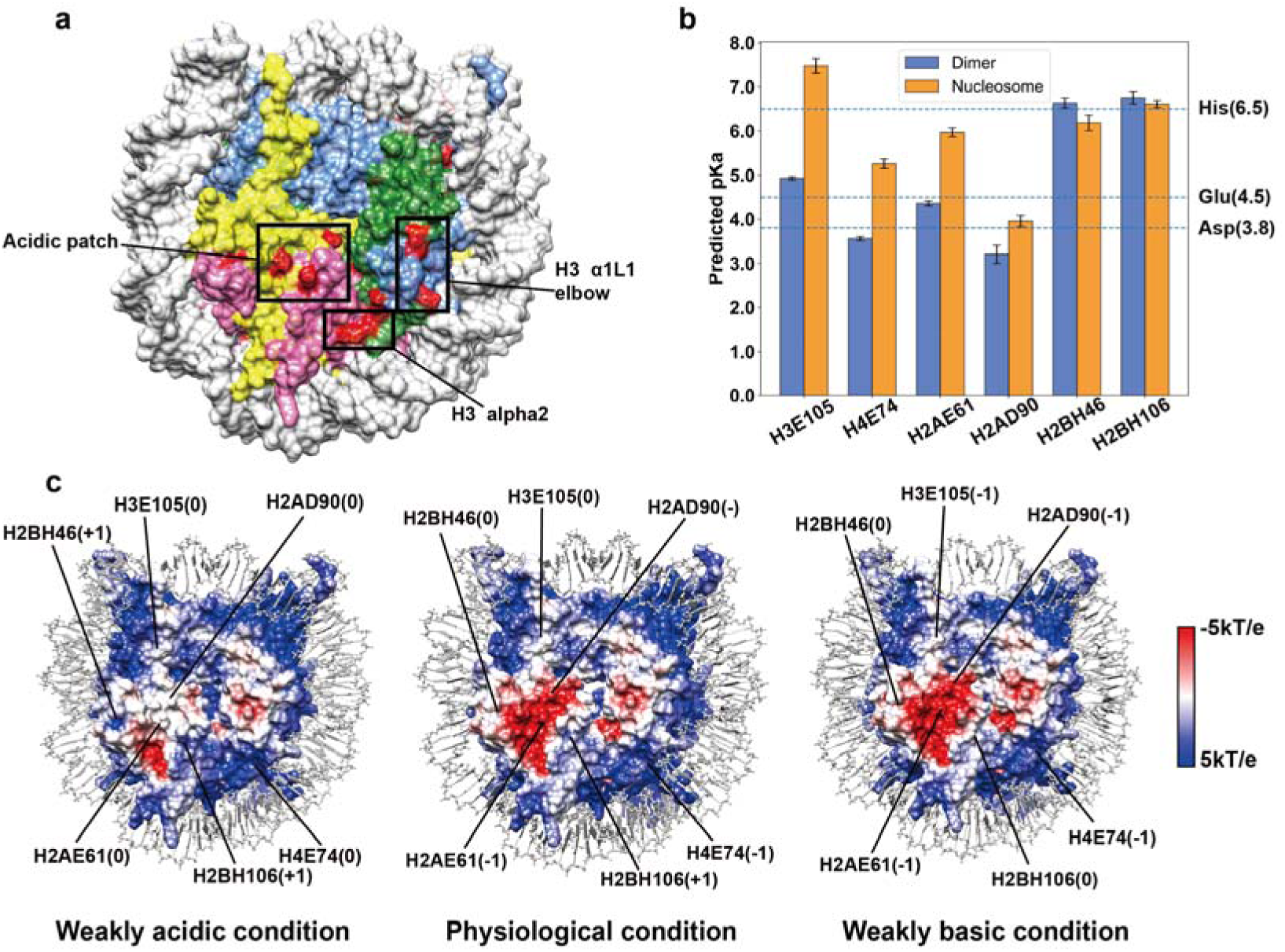
Modulation of nucleosome surface electrostatic potentials by protonation states of histone ionizable residues. (a) Locations of identified histone titratable residues whose protonation states are sensitive to pH perturbations in cellular conditions. (b) Six histone titratable residues are located on the surface of the histone octamer, whose protonation states are sensitive to pH perturbations. (c) Nucleosome surface electrostatic potentials under different pH conditions: Left: under weakly acidic condition (pH 5-6.5); Middle: within the physiological pH range (pH 6.5-7.5); Right: under weakly basic condition (pH 7.5-9). The electrostatic potential is mapped onto the surface of histone octamer on a scale from +5 kT/e (blue) to −5 kT/e (red).

### Protonation states of key histone residues modulate nucleosome surface electrostatic potential

We have demonstrated that histone residues within nucleosomes undergo substantial pKa shifts due to their protein environments. Such shifts in pKa values can indeed correspond to changes in the protonation states of histone ionizable residues, leading to alterations in nucleosome charges. Based on our pKa predictions, we categorized histone titratable residues into four classes based on their protonation states at different pH levels (see methods section and Supplementary Table 7). We identified 14 histone titratable residues whose protonation states were sensitive to changes in pH within the typical cellular environment (Supplementary Table 8). Additionally, four histone titratable residues exhibited protonation state changes even within physiological pH condition (pH 6.5-7.5), including H2BGlu71, H2BHis106, H3Glu97, and H3Glu105. According to the calculated pKa values, all four residues were protonated within physiological pH range from 6.5 to 7.5. H2BHis106 carried a positive net charge, while the protonated forms of H2BGlu71, H3Glu97, and H3Glu105 had neutral charges.

Since alterations in the protonation states of histone residues can influence the surface charges of the histone octamer, we investigated how these changes can affect nucleosome surface electrostatic potentials. Among the 14 identified histone titratable residues whose protonation states are sensitive to pH changes in cellular conditions, we found six located on the surface of the histone octamer, including H2AE61, H2AD90, H2BH46, H2BH106, H3E105 and H4E74, mostly located around acidic patch, H3 Alpha2-helix and H4 Alpha2-helix (Figure 3). Changes in their protonation states can significantly impact the nucleosome surface electrostatic potentials.

Moreover, we performed electrostatic analysis of nucleosome structures with histone titratable residues in different pH environments (Figure 3c). Interestingly, we observed significant alterations in nucleosome surface electrostatic potentials resulting from changes in the protonation states of key histone residues. Specifically, under weakly acidic condition (pH 5-6.5), the protonation of H2AGlu61, H2AD90, H2BHis46, H4Glu74 and H2BH106 led to a substantial shift from negative electrostatic potentials to neutral or positive potentials near the acidic patch and H4 Alpha2-helix. Conversely, the deprotonation of H2BHis106 resulted in enhanced negative potentials under slightly basic conditions (pH 7.5-9). Overall, our results highlight that alterations in the protonation states of several key histone residues can occur due to perturbations in pH near cellular physiological conditions, leading to significant effects on nucleosome surface electrostatic potentials and potentially impacting nucleosome interactions and higher-order chromatin structures.

### Histone residue protonation states regulate the long-range electrostatic interactions between nucleosome and chromatin factors

Changes in the protonation states of histone residues can lead to modifications in the electrostatic potential of the nucleosome surface. These alterations have the potential to influence interactions between nucleosomes and chromatin factors. To confirm our hypotheses, we conducted systematic analyses of the long-range electrostatic forces generated between nucleosomes and different binding proteins using representative nucleosome complex structures. These analyzed complexes contain seven distinct nucleosome-binding proteins known for their critical roles in processes such as PTM reading and writing, chromatin remodeling, DNA repair, and DNA replication (Supplementary Table 9).

Our findings demonstrate that under cellular pH conditions, the strengths and directions of electrostatic forces can be significantly affected by changes in the protonation states of key histone titratable residues (Figure 4). For example, we observed significant changes in the magnitude and direction of the electrostatic interaction between the nucleosome and Tumor suppressor p53-binding protein 1 across different pH ranges (Figure 4 and Supplementary Figure 2). Specifically, under weakly acidic conditions (pH 5 to 6.5), the strength of the electrostatic force between the nucleosome and binding protein is minimal, while it reaches its maximum values under weakly basic conditions (pH 7.5 to 9). Moreover, the directions of the electrostatic forces are also shifted, indicating weakened electrostatic interactions between the nucleosome and binding proteins under weakly acidic conditions. Such trends were also observed for other nucleosome binding proteins such as PRC1, RSC, DOT1L and SIR3 (Figure 4 and Supplementary Figures 2 and 3)

**Figure 4.**
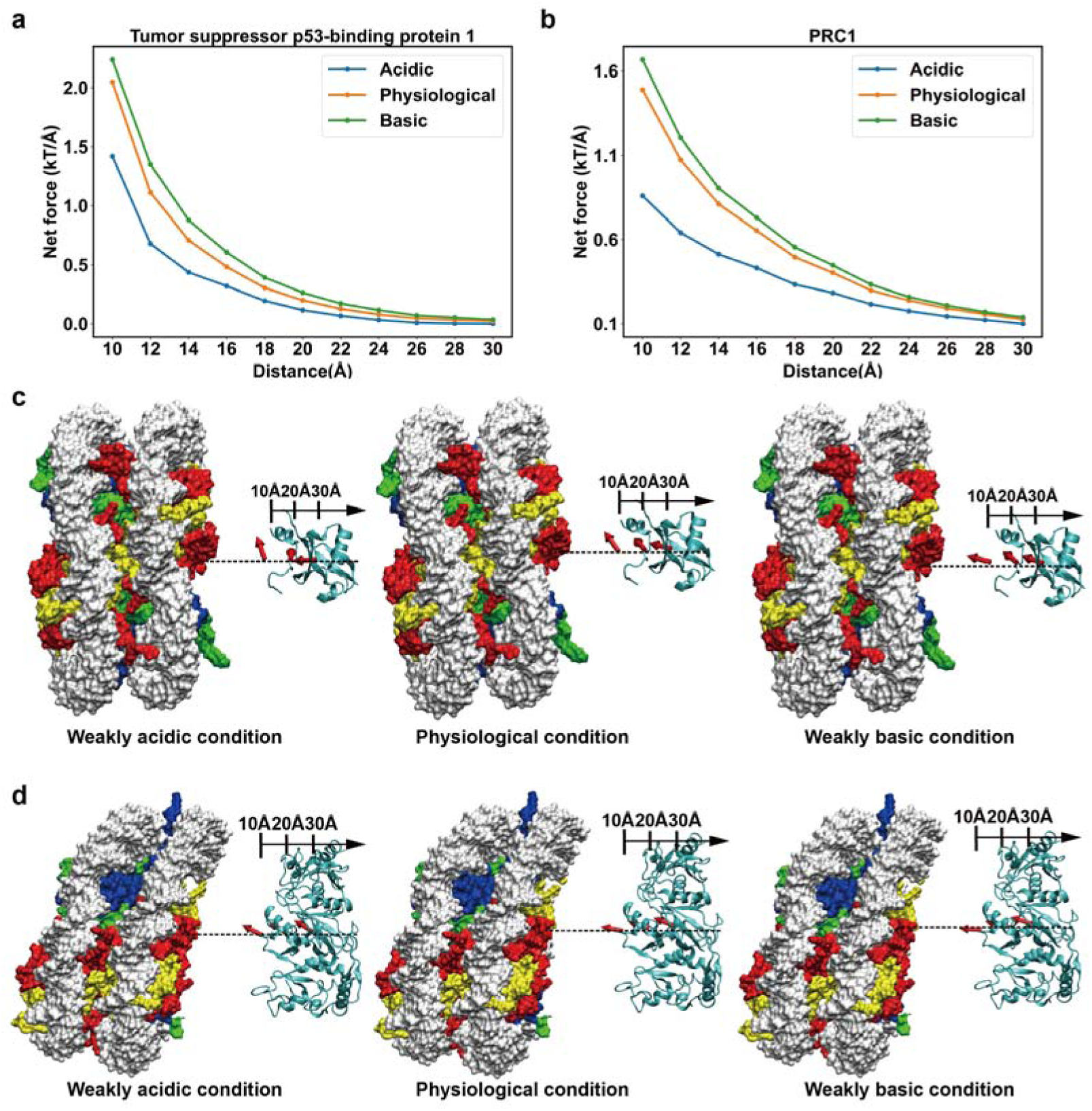
Protonation states of histone ionizable residues regulate the long-range electrostatic interactions between nucleosomes and regulatory proteins. (a) and (b) Comparison of the magnitudes of electrostatic forces between nucleosomes and binding proteins (Tumor suppressor p53-binding protein 1 on the left, and PCR1 on the right) under weakly acidic (pH 5 to 6.5), physiological (pH 6.5 to 7.5), and weakly basic (pH 7.5 to 9) conditions. The electrostatic forces were computed at various center-of-mass separation distances ranging from 10 to 30 Å, with 2 Å increments, using PDB structures 5KGF and 8GRM. (c) and (d) Comparison of the directions of electrostatic forces between nucleosomes and binding proteins (Tumor suppressor p53-binding protein 1 above, and PCR1 below) under weakly acidic (pH 5 to 6.5), physiological (pH 6.5 to 7.5), and weakly basic (pH 7.5 to 9) conditions. The directions of electrostatic interactions are represented by red arrows at separating distances of 10 Å, 20 Å, and 30 Å.

For the majority of analyzed nucleosome complexes, we observed that nucleosomes generally exhibited the strongest long-range electrostatic interactions with binding proteins under physiological or weakly basic conditions (Figure 4 and Supplementary Figure 3). This observation underscores the pivotal role of protonation state changes of key histone residues in modulating nucleosome surface electrostatic potentials. Specifically, under physiological or slightly basic pH conditions, nucleosome can maintain a stronger negative electrostatic potential around the acidic patch, H3 Alpha2-helix and H4 Alpha2-helix, thereby enhancing electrostatic interactions between the histone octamer and binding proteins.

### Binding of regulatory proteins to nucleosome is coupled with proton uptake or release of histone titratable residues

Nucleosome harbors a plethora of interactions with various chromatin factors, and their binding interfaces are enriched with ionizable groups. Consequently, it is reasonable to expect that proton uptake or release of these ionizable groups occurs upon the binding of proteins to nucleosomes. However, the underlying mechanisms of this phenomenon have not been systematically investigated. To address this gap, we conducted a systematic analysis using 83 representative nucleosome complex structures. These structures encompass nucleosome interactions with 200 distinct binding proteins, essential for a wide array of epigenetic regulation processes such as PTM reading, writing, and erasing, DNA repair and replication, chromatin remodeling, and gene transcription (Supplementary Figure 1).

For each nucleosome complex structure, we calculated pKa shifts and analyzed protonation state changes of histone residues resulting from the binding of regulatory proteins to nucleosomes (see the Methods section for details). Our analyses revealed positive pKa shifts of acidic residues and negative shifts of basic residues upon regulatory protein binding to nucleosomes, indicating suppression of acidity and basicity of histone ionizable groups (Figure 5a). Furthermore, we identified 14 titratable acidic residues and 19 titratable basic residues exhibiting significant pKa shifts upon protein binding to nucleosomes (|ΔpKa| ≥ 1.5, observed in at least three different complex structures), with the largest changes up to 5.9 (Figure 5a). Among these residues, three are susceptible to proton uptake or release within physiological pH condition (H3E74, H2AE65, and H2AE93) and eight are susceptible to weakly acidic or basic pH conditions (Supplementary Table 10). These residues are distributed among all four histone types and are mostly located around the acidic patch and H3α1L1 elbow, suggesting potential effects on nucleosome surface electrostatics and interactions with partner proteins (Figure 5c).

**Figure 5.**
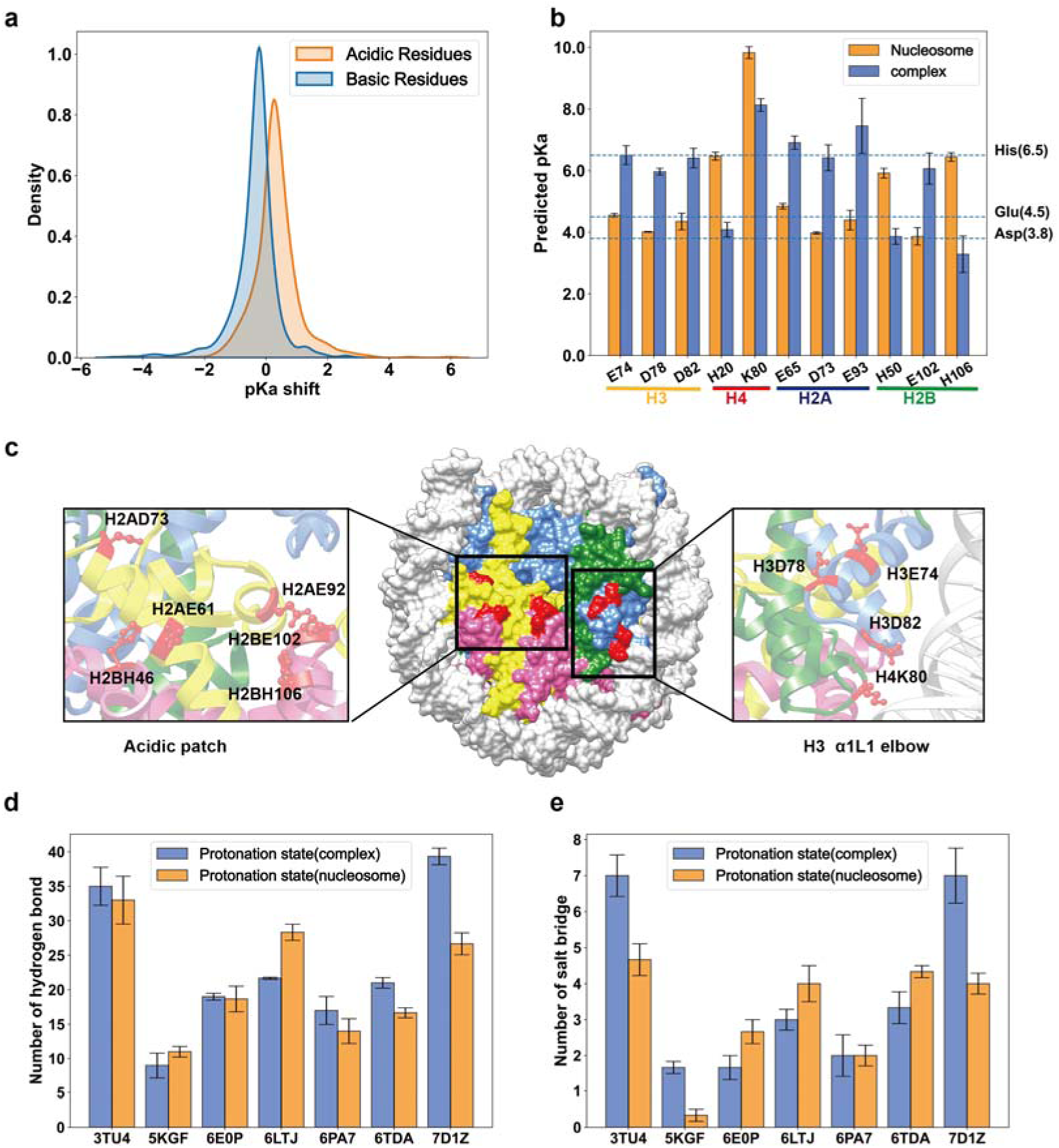
Effects of proton uptake or release of histone titratable residues on nucleosome-chromatin factor interactions. (a) Distribution of pKa shifts of histone titratable residues caused by the interactions between nucleosomes and binding proteins. The pKa shift represents the change in pKa value of the titratable group from the unbound state to the bound state. (b) and (c) Histone titratable residues that exhibited protonation state change upon binding of partner proteins to nucleosome. These residues were mapped onto the surface of nucleosome structure using the PDB 2PYO (shown as red). Protein H2A, H2B, H3 and H4 were shown as pink, yellow, green, and blue, respectively. (d) and (e) Comparison of the hydrogen bonds and salt bridges formed between nucleosomes and binding partner proteins under two different protonation states of histone titratable residues: in one set, histone residues were assigned protonation states corresponding to the unbound condition, while in the other set, histone residues were assigned protonation states reflecting the bound condition.

Next, we investigated how such proton uptake or release of histone titratable residues could affect nucleosome-partner protein interactions. To answer this question, we performed molecular dynamics simulations of seven representative nucleosome complex structures by setting the protonation states of histone residues in two states: one with the protonation states when nucleosome bound with chromatin factors and another with the protonation states in the unbound state. Our calculations indicated that proton uptake or release in histone residues can generally promote the binding of proteins to nucleosome structures and increase the binding free energies (Supplementary Table 11). Indeed, in six out of the seven cases, we observed an increase in nucleosome-partner binding free energies with a mean value of 11.09 kcal/mol. Furthermore, we analyzed the hydrogen bonds and salt bridges formed between histones and binding proteins under different protonation states. Despite the decreased number of salt bridges due to changes in histone ionizable group protonation states in several cases, we observed a significant increase in hydrogen bonds within the binding interfaces, thereby promoting nucleosome-partner protein interactions (Figure 5).

### Histone cancer mutations alter nucleosome surface electrostatic potential and the long-range electrostatic interactions

Histone cancer mutations predominantly cluster on the histone binding interfaces, exerting profound disruptive effects on nucleosome interactions. Leveraging our recently compiled comprehensive dataset of histone cancer mutations (see method section), we systematically examined the impact of recurrent cancer mutations on nucleosome surface charges and electrostatics. Our analysis revealed that over 57% of recurrent histone cancer mutations altered the net charges on the surface of the histone octamer, with more than 20% of the mutations even flipping residue charges (Figure 6). Notably, approximately 58% of these charge-altering mutations are situated on acidic patches, H3α1L1 elbow, H3α2, and H3 tail regions (histone H1 cancer mutations were excluded in analysis, Supplementary Figure 4).

**Figure 6.**
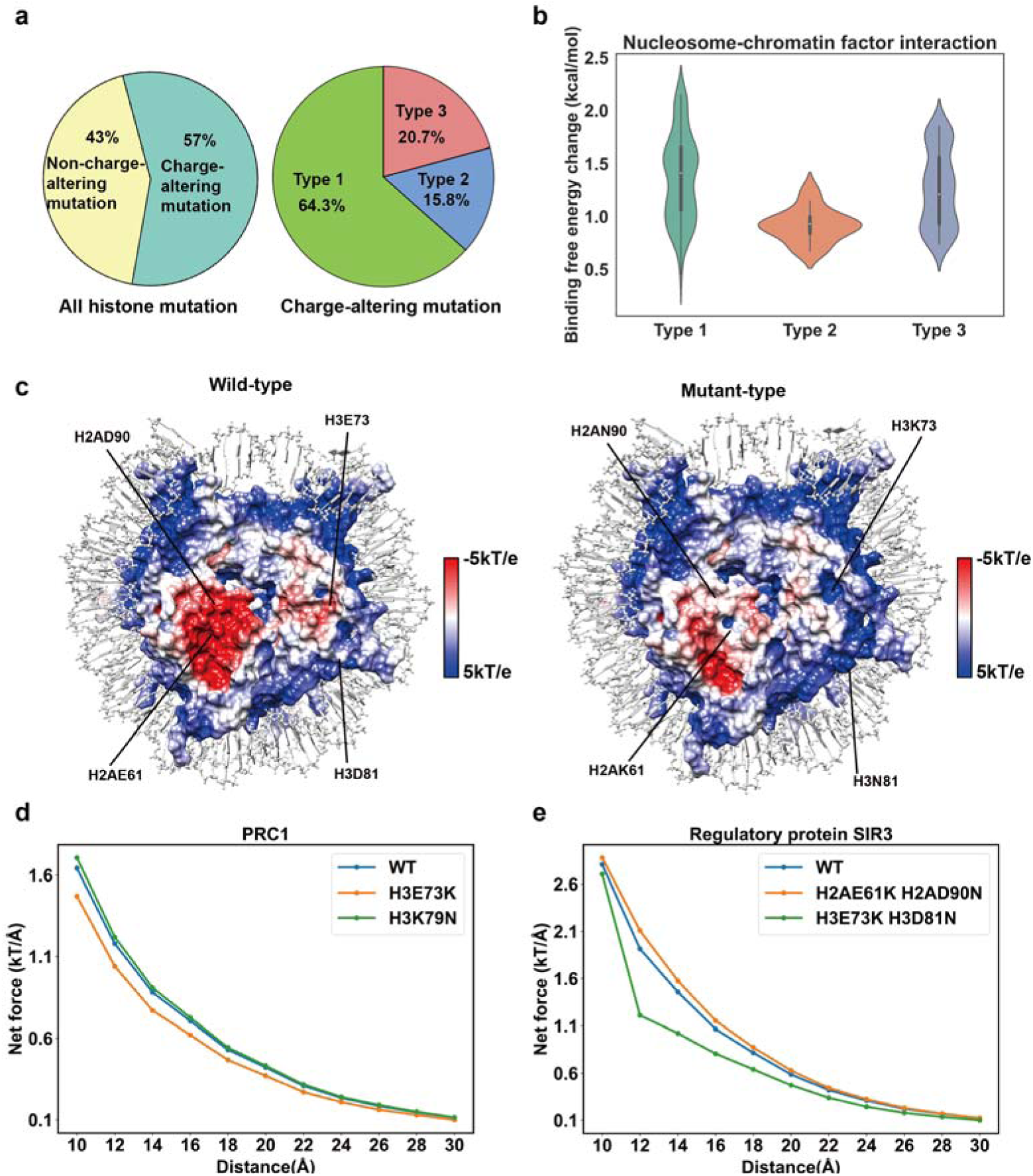
Impact of histone cancer mutations on nucleosome surface electrostatic potential and electrostatic interactions. (a) Classification of recurrent histone cancer mutations based on their impact on residue net charges. Among the charge-altering mutations, Type 1 represents a charged residue mutated to a neutral residue, Type 2 represents a neutral residue mutated to a charged residue, and Type 3 represents mutations that flip the net charge of residues. (b) Distribution of binding free energy changes (ΔΔGs) caused by recurrent histone cancer mutations in nucleosome-partner protein interactions for different mutation types. (c) Changes in the nucleosome surface electrostatic potential caused by histone cancer mutations: left, wild-type; right, mutant-type. The electrostatic potential is mapped onto the surface of the histone octamer on a scale from +5 kT/e (blue) to −5 kT/e (red). (d) and (e) Comparison of the magnitudes of electrostatic forces between the nucleosome and binding partners in wild-type and mutant states. The electrostatic forces were computed at various center-of-mass separating distances ranging from 10 to 30 Å, with 2 Å increments, using PDB structures 8GRM and 3TU4.

To investigate the consequences of charge-altering mutations on nucleosome-partner binding energies, we calculated the binding free energy changes (ΔΔGs) of 153 histone recurrent mutations that were located on the binding interfaces of nucleosome-partner complex structures and altered the residue net charge. Our calculations revealed the strong destabilizing effects of these mutations on nucleosome-partner protein binding, with a mean decrease in binding free energy values of 1.29 kcal/mol, especially for the Type 1 and Type 3 mutations (Figure 6).

Through electrostatic potential calculations, we demonstrated that even one or two histone cancer mutations can induce remarkable effects on the nucleosome surface electrostatic potentials (Figure 6 and Supplementary Table 12), with the mutations H2AE61K and H3E73K exhibiting particularly pronounced effects. Furthermore, we assessed the impact of charge-altering mutations on electrostatic interactions using nucleosome complex structures (Supplementary Table 13) and our results revealed their disruptive effects on the long-range electrostatic interactions between nucleosomes and various chromatin factors, such as PRC1, Regulatory protein SIR3 and Centromere proteins (Figure 6 and Supplementary Figure 5). We show that the strengths and directions of electrostatic forces between nucleosome and binding proteins can be significantly altered by even a single histone cancer mutation. These impacts on nucleosome-partner protein binding affinities and long-range electrostatic interactions are expected to have profound effects on chromatin structures and various epigenetic regulation processes.

### Histone residue protonation state and cancer mutation impact nucleosome stacking in higher-order structures

It is certain that the electrostatic interactions of nucleosomal particles can play a crucial role in chromatin folding processes and in mediating nucleosome self-association^41–44^. For instance, the acetylation of histone tails reduces their positive charge, thereby disrupting the electrostatic interactions among nucleosomal particles and leading to less compact chromatin structures^13, 21, 45, 46^. In our study, we have demonstrated that changes in histone residue protonation or cancer mutations can alter both nucleosome surface electrostatic potentials and long-range electrostatic interactions, potentially influencing nucleosome self-association. To validate our hypothesis, we have further performed electrostatic analyses using two experimentally determined nucleosome higher-order structures. One is a structure of an H1-bound 6-nucleosome array (PDB: 6HKT)^47^, while another is one telomeric tetranucleosome structure (PDB: 7V9K)^48^.

Our findings reveal that changes in the protonation states of histone ionizable residues can significantly impact nucleosome-nucleosome electrostatic interactions under cellular pH conditions (Figure 7). In different neighboring nucleosomes, the electrostatic forces can be either repulsive or attractive depending on the orientations of nucleosomes in stacking. Additionally, the strengths and directions of electrostatic forces between neighboring nucleosomes can be notably affected by alterations in the protonation states of key histone titratable residues. Specifically, we observed that nucleosomes generally exhibited the strongest attractive or the weakest repulsive electrostatic forces by neighboring nucleosomes under weakly acidic conditions, thereby promoting nucleosome stacking and compaction.

**Figure 7.**
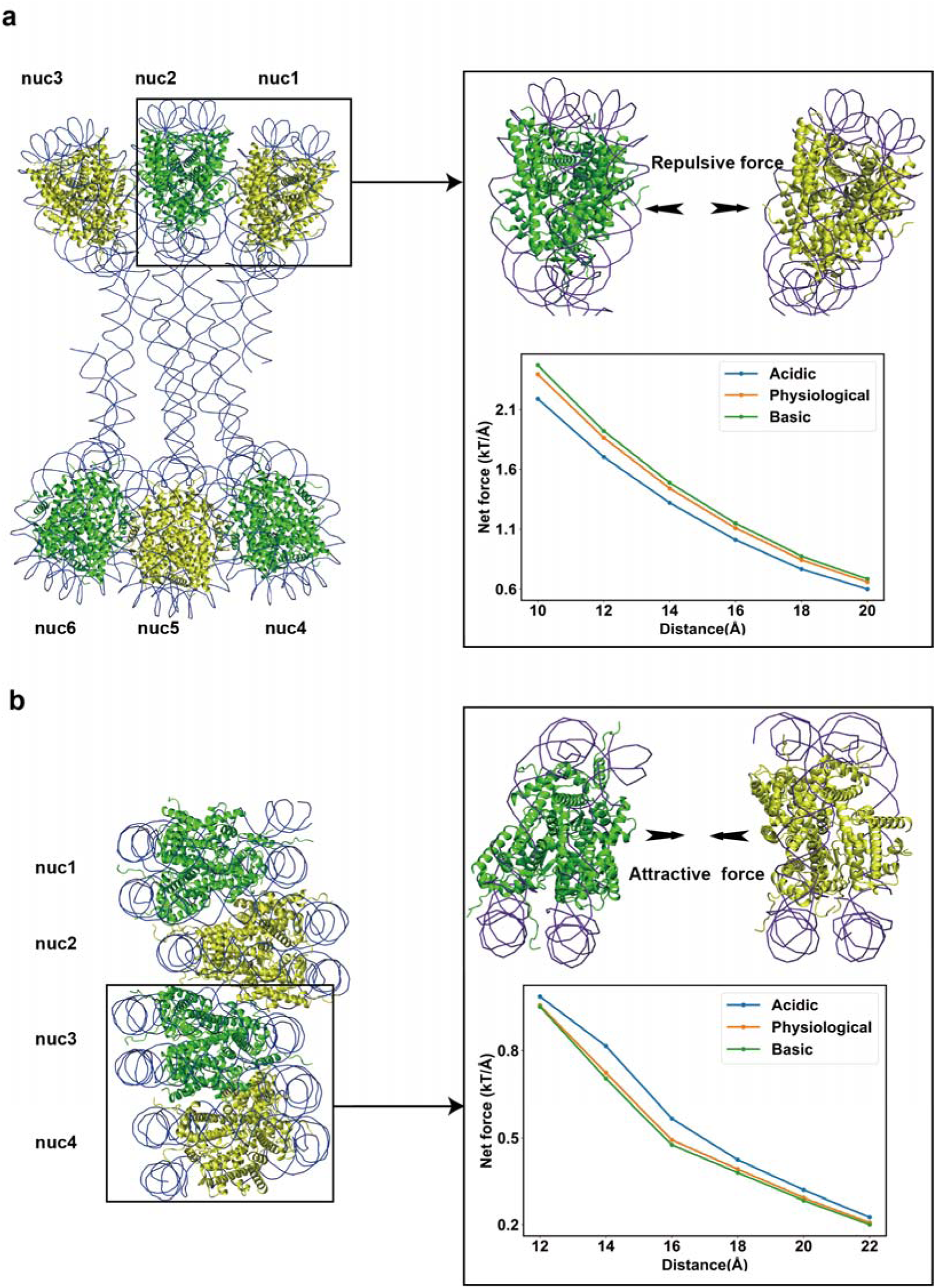
Protonation states of histone titratable residues regulate the long-range electrostatic interactions in higher-order nucleosome structures. (a) and (b) The strengths and directions of electrostatic forces between neighboring nucleosomes in two structures of nucleosome arrays: H1-bound 6-nucleosome array (top) and telomeric tetranucleosome structure (bottom). The electrostatic forces were computed at various center-of-mass separating distances using PDB structures 6HKT and 7V9K under weakly acidic (pH 5 to 6.5), physiological (pH 6.5 to 7.5), and weakly basic (pH 7.5 to 9) conditions.

In addition to nucleosome-nucleosome interactions, we also investigated the impact of cancer mutations or pH perturbations on the binding between the H4 tail and the acidic patch. Our results indicated that the long-range electrostatic interactions can be considerably affected by the pH perturbations even in the cellular conditions (Figure 8 and Supplementary Figure 7). The H4 tail-acidic patch interactions would be dramatically weakened when the pH was shifted to slightly acidic condition (pH from 5 to 6.5), thereby modulating the nucleosome compaction. Moreover, we identified six charge-altering cancer mutations on H4 tails and on the acidic patches, potentially leading to significantly effects on the electrostatic interactions (Supplementary Table 14). Using a model of the H4 tail-nucleosome complex built with AlphaFold3^49^, we calculated the electrostatic forces between the H4 tail and the acidic patch in wild-type and mutant states. Our results show that even a single charge-altering cancer mutation can lead to substantial changes in electrostatic interactions between the H4 tail and an adjacent nucleosome, thereby affecting the compaction of nucleosome arrays (Figure 8 and Supplementary Figure 7). Overall, our findings suggest that alterations in histone protonation or cancer mutations may not only impact individual nucleosomes but also influence their collective behavior, such as self-association, thus contributing to the overall organization and dynamics of chromatin structure.

**Figure 8.**
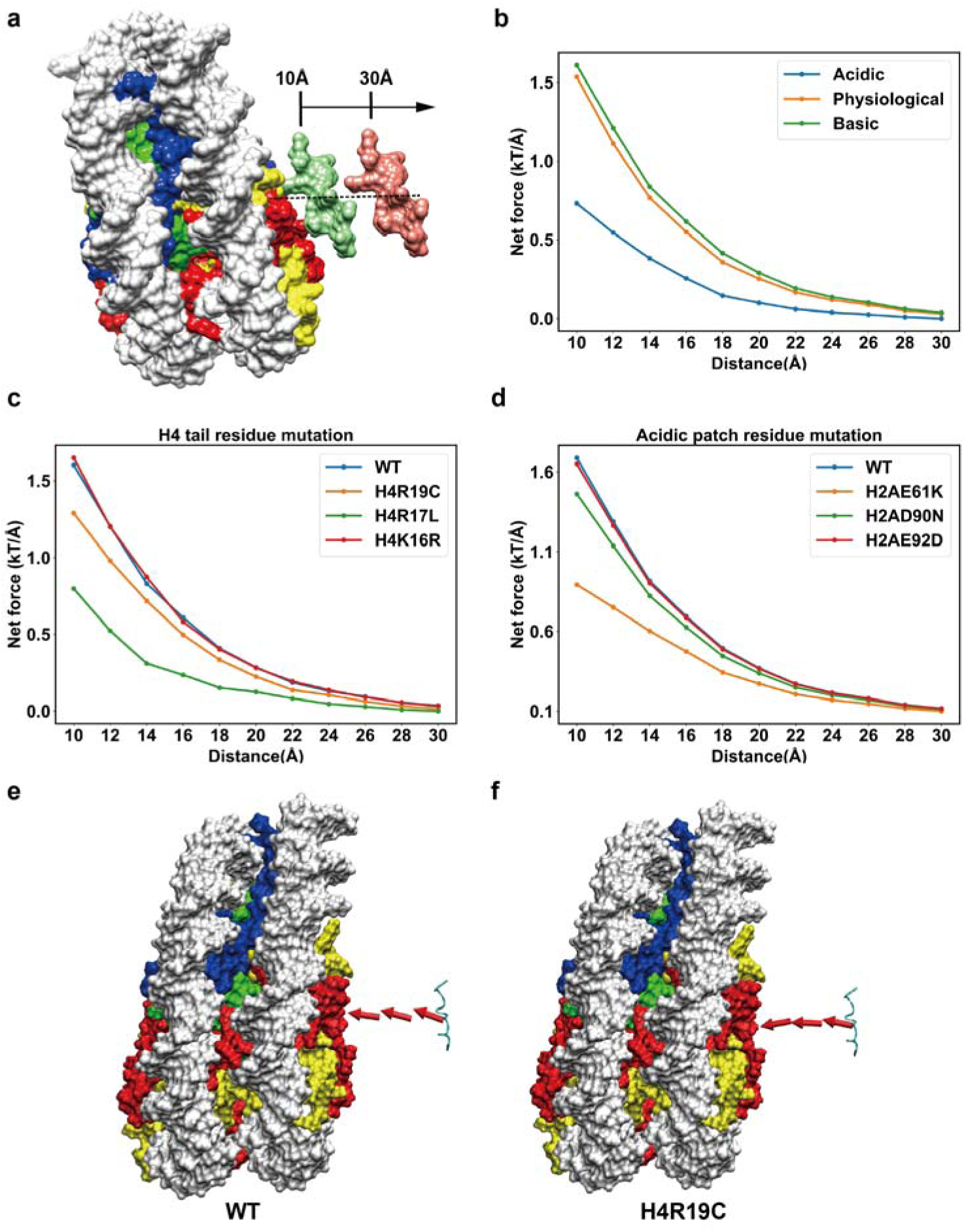
Histone cancer mutations affect the electrostatic interactions between the H4 tail and acidic patch. (a) Analyses of the long-range electrostatic forces generated between nucleosomes and H4 tail at separating distances from 10 to 30 Å, along the line connecting their centers of mass. (b) Comparison of the magnitudes of electrostatic forces between nucleosome and histone H4 tail under weakly acidic (pH 5 to 6.5), physiological (pH 6.5 to 7.5), and weakly basic (pH 7.5 to 9) conditions. (c) and (d) Comparison of the magnitudes of electrostatic forces between the H4 tail and acidic patch in wild-type and mutant states. The electrostatic forces were computed at various center-of-mass separating distances ranging from 10 to 30 Å, with 2 Å increments. (e) and (f) Comparison of the directions of electrostatic forces between nucleosomes and H4 tails in wild-type and mutant states. The directions of electrostatic interactions are represented by red arrows at separating distances of 10 Å, 20 Å, and 30 Å.

## Discussion

Electrostatic interactions are pivotal in biomolecular interactions, acting as long-range forces that govern the protein-protein and protein-nucleic acid interactions ^50–55^. While nucleosome formation reduces the net charge of DNA within chromatin, the nucleosome itself still maintains a strong negative electrostatic field^12^. Recent studies have emphasized the importance of electrostatic interactions in mediating nucleosome interactions with various chromatin factors, owing to the complementary charges present in the histone octamer and partner protein binding interfaces^5, 20, 56–59^. We show that histone residues in nucleosomes undergo significant pKa changes due to their protein environments, characterized by elevated pKa values for histone acidic residues and decreased values for basic residues. Our findings demonstrate that even minor perturbations in pH near cellular physiological conditions can profoundly alter nucleosome surface electrostatics, particularly in regions surrounding acidic patches. Elevated pKa values result in protonation of H4E74, H2AE61, H3E105 and H2AD90 within slightly acidic pH conditions, significantly dampening negative charges. Additionally, H2BH106 and H2BH46 become significantly protonated at slightly acidic pH, further reducing the electronegativity and hydrogen bonding capacity of the acidic patch. Utilizing experimentally determined nucleosome-partner protein complex structures, we have demonstrated that changes in the protonation states of key histone residues, such as H2AE93, H2BE103 and H3D82, can modulate the binding processes of various chromatin factors to nucleosomes by altering the directions and magnitudes of long-range electrostatic forces. Our findings indicate that nucleosomes typically exhibit stronger electrostatic forces with partner proteins under physiological or slightly basic pH conditions, whereas these forces are notably diminished under slightly acidic conditions.

In our investigation, we observed significant changes in pKa values and protonation states of ionizable groups accompanying nucleosome-partner protein binding, consistent with previous findings^16–18, 60^. Specifically, we identified 14 titratable acidic residues and 19 titratable basic residues exhibiting notable pKa shifts upon protein binding to nucleosomes, indicative of proton uptake or release during binding processes. Furthermore, our analyses revealed that the proton uptake or release of histone titratable residues generally promoted the formation of hydrogen bonds in the histone-partner protein binding interfaces, consequently enhancing their binding affinities. These findings underscore the crucial roles of protonation states of ionizable groups in modulating nucleosome-partner protein interactions, emphasizing the importance of considering these factors in future studies.

Electrostatic interactions serve as the primary driving forces in the folding of arrays of nucleosomes into chromatin^8, 9^. Notably, recent experiments have highlighted the pivotal role of electrostatic interactions between nucleosomes as a major biophysical driving force for organizing nucleosomes into euchromatin and heterochromatin^61^. Furthermore, experimental evidence has demonstrated that charge-altering mutations at nucleosome-nucleosome interaction interfaces significantly impact chromatin compaction^62–65^. Utilizing two experimentally determined structures of nucleosome arrays, we demonstrate that even slight perturbations in pH conditions can induce significant alterations in nucleosome-nucleosome and H4 tail-acidic patch electrostatic interactions. Additionally, leveraging our previously constructed histone cancer mutation dataset, we show that even a single cancer mutation can induce significant alterations in the electrostatic interactions between H4 tail and acidic patch of neighboring nucleosome.

We conclude that electrostatic interactions are crucial in orchestrating the binding and recognition of various chromatin factors to nucleosomes and the folding of nucleosome arrays into chromatin (Figure 9). Histone ionizable residues are highly sensitive to cellular pH fluctuations, leading to changes in their protonation states and consequent alterations in nucleosome surface electrostatic potentials and interactions. Such changes in protonation states should not be overlooked in studies of nucleosome-partner interactions and may contribute to intrinsic regulatory mechanisms. Additionally, we propose that alterations in histone residue protonation or cancer mutations have the potential to impact nucleosome self-association and higher-order chromatin structures. However, quantitatively characterizing these dynamic processes remains challenging. Future research efforts should focus on developing experimental and computational techniques to elucidate the spatial and temporal hierarchy of dynamic chromatin processes, thereby bridging the gap in our understanding.

**Figure 9.**
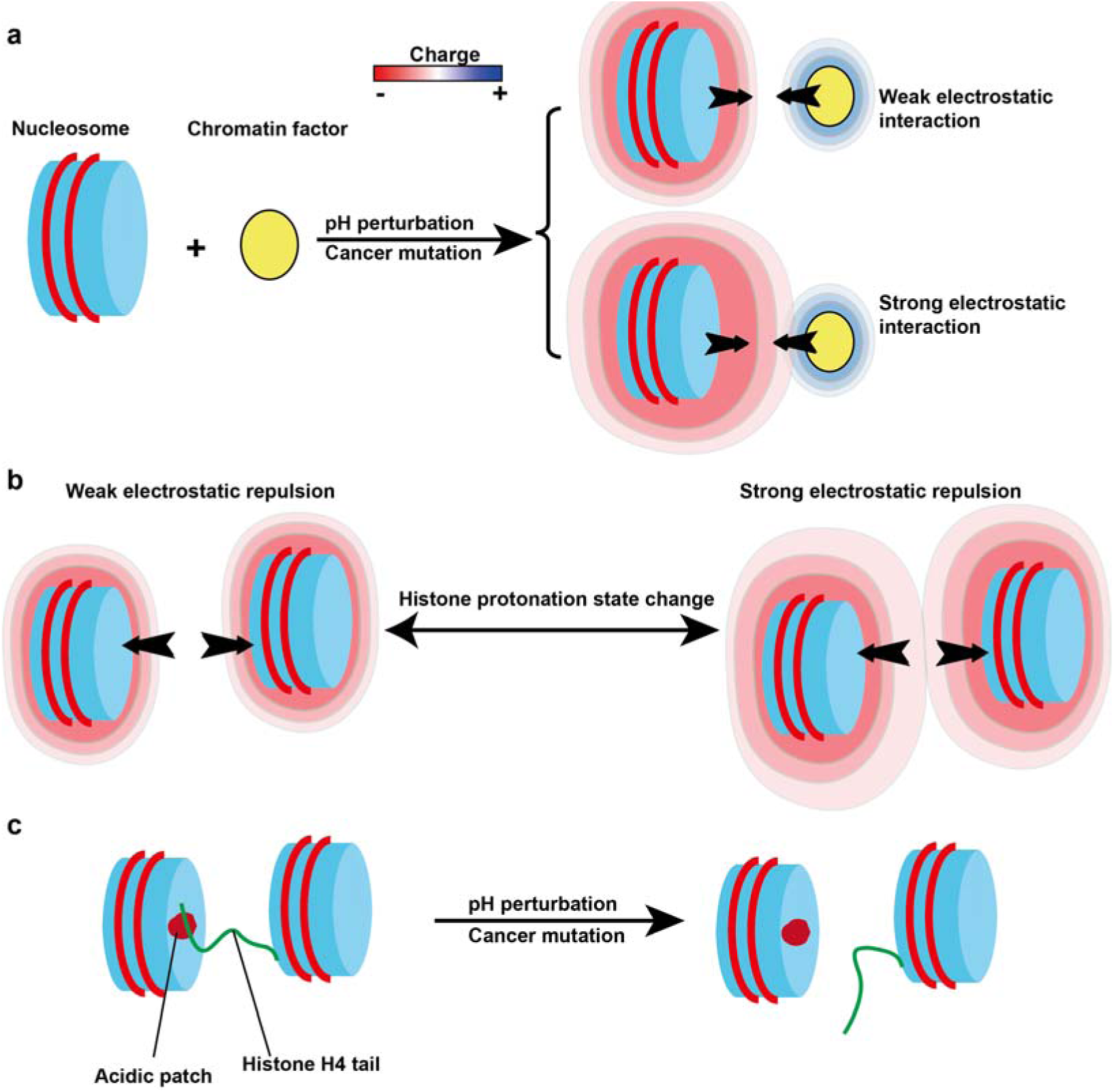
A generalized model explaining pH perturbation and cancer mutation can affect nucleosomes’ interactions with chromatin factors and higher-order nucleosome structures by modulating the long-range electrostatic interactions. (a) Perturbations within the cellular pH ranges and charge-altering cancer mutations can affect the interactions between nucleosomes and chromatin factors. (b) Protonation states of histone ionizable residues can regulate nucleosome stacking in higher-order nucleosome structures. (c) Perturbations within the cellular pH ranges and charge-altering cancer mutations can affect the interactions of H4 tail with acidic patch in neighboring nucleosome.

## Acknowledgments

This work was supported by the National Natural Science Foundation of China (No.12205112) and Fundamental Research Funds for Central China Normal University. LJJ was supported by National Nature Science Foundation of China (No. 22377029) and Fundamental Research Funds for Central China Normal University.

