## Supplementary Information for "Electrostatic interactions in nucleosome and higher-order structures are regulated by protonation state of histone ionizable residue"

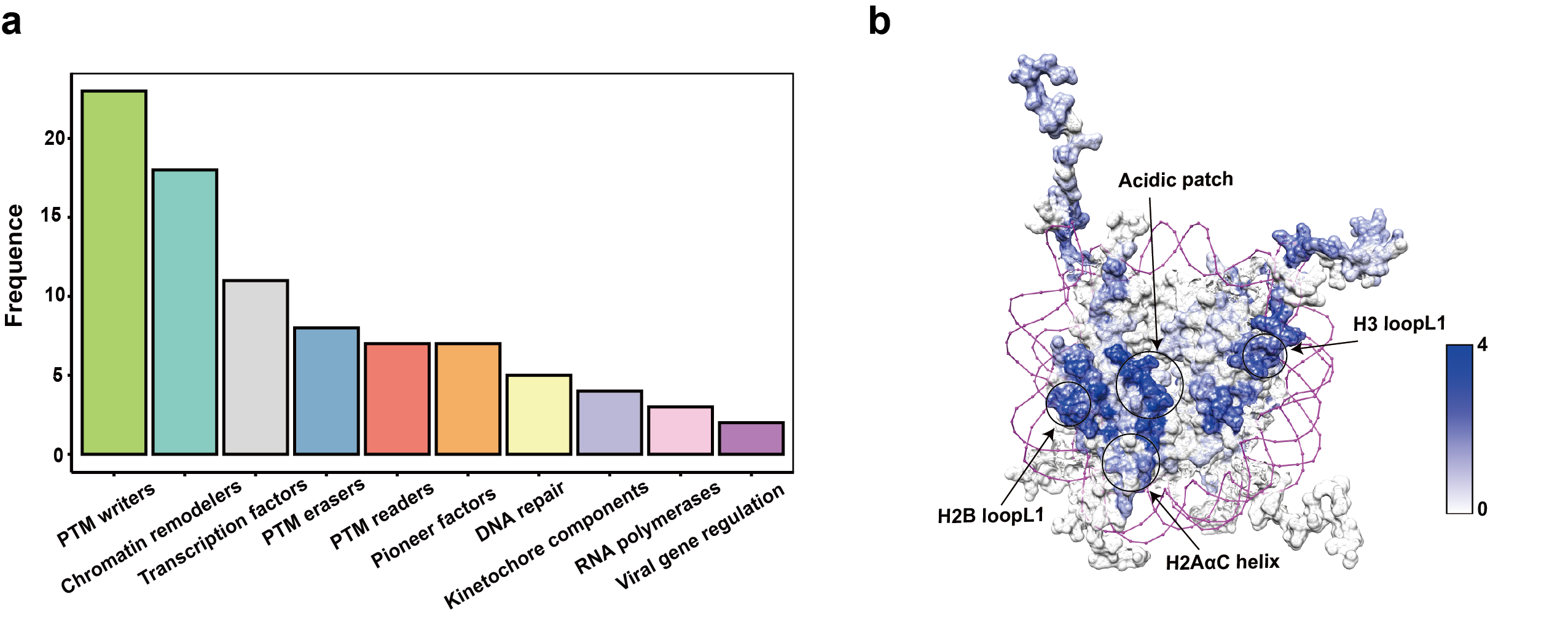

**Supplementary Figure 1.** (a) Functional classification of 83 representative nucleosome complexes. (b) Mapping of binding interfaces of chromatin factors onto the histone octamer. The colors indicate the number of binding proteins per residue on the surface of the octamer.

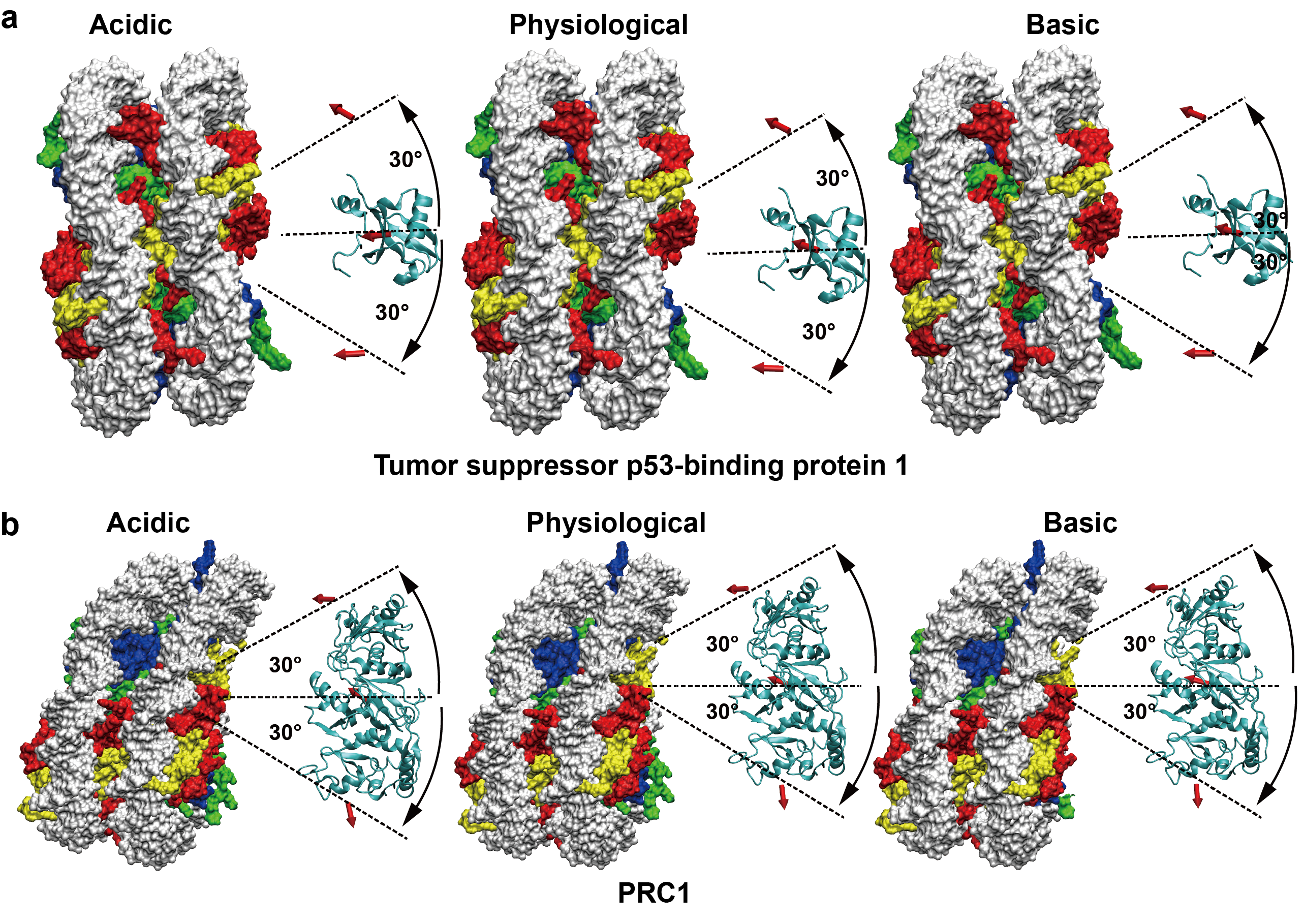

**Supplementary Figure 2.** The electrostatic forces between nucleosomes and regulatory proteins at different rotation angles of the line connecting their centers of mass. (a) and (b) Comparison of the directions of electrostatic forces between nucleosomes and binding proteins (Tumor suppressor p53-binding protein 1 above (PDB:5KGF), and PCR1 below (PDB:8GRM)) under weakly acidic (pH 5 to 6.5), physiological (pH 6.5 to 7.5), and weakly basic (pH 7.5 to 9) conditions. The directions of the electrostatic forces are represented by red arrows at rotation angles ranging from -30˚ to 30˚.

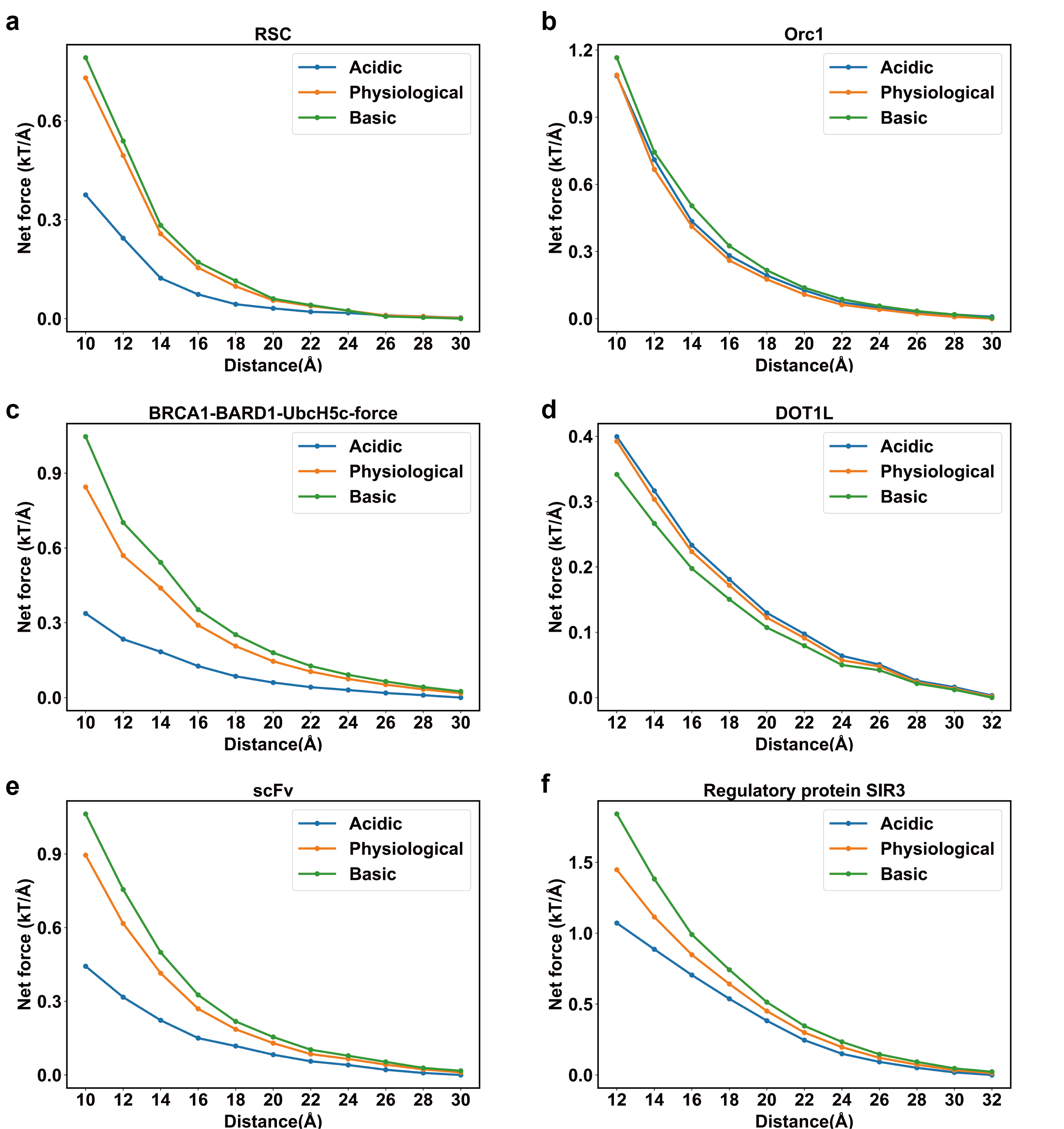

**Supplementary Figure 3.** Comparison of the magnitudes of electrostatic forces between nucleosomes and different binding proteins under weakly acidic (pH 5 to 6.5), physiological (pH 6.5 to 7.5), and weakly basic (pH 7.5 to 9) conditions. The electrostatic forces were computed at various center-of-mass separating distances using PDB structures 6TDA, 7E9F, 7LYB, 7XCT, 6E0P, and 3TU4.

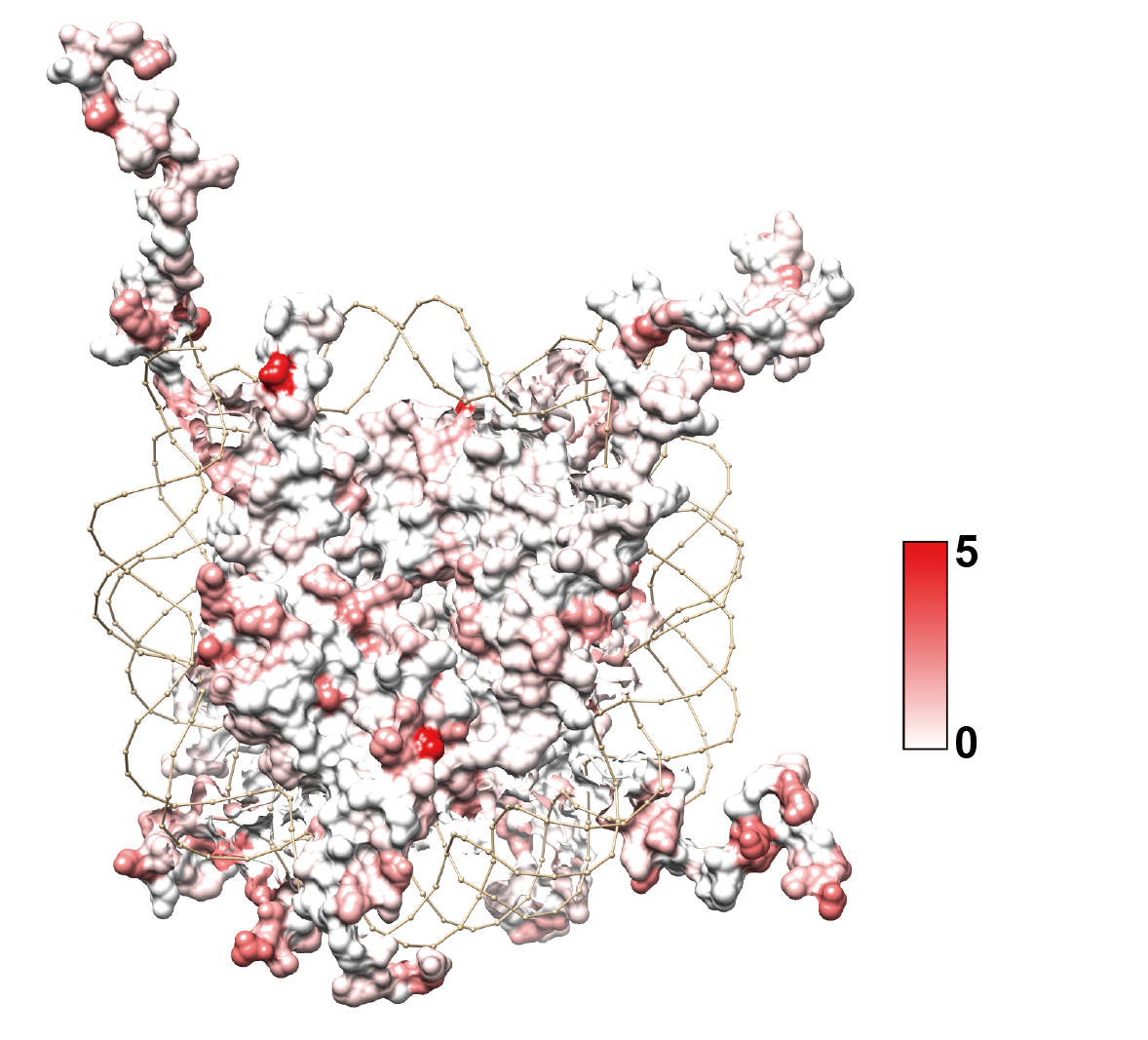

**Supplementary Figure 4.** Mapping of recurrent histone cancer mutations onto the surface of the histone octamer within the nucleosome. The number of mutations per residue is indicated by intensities of red color.

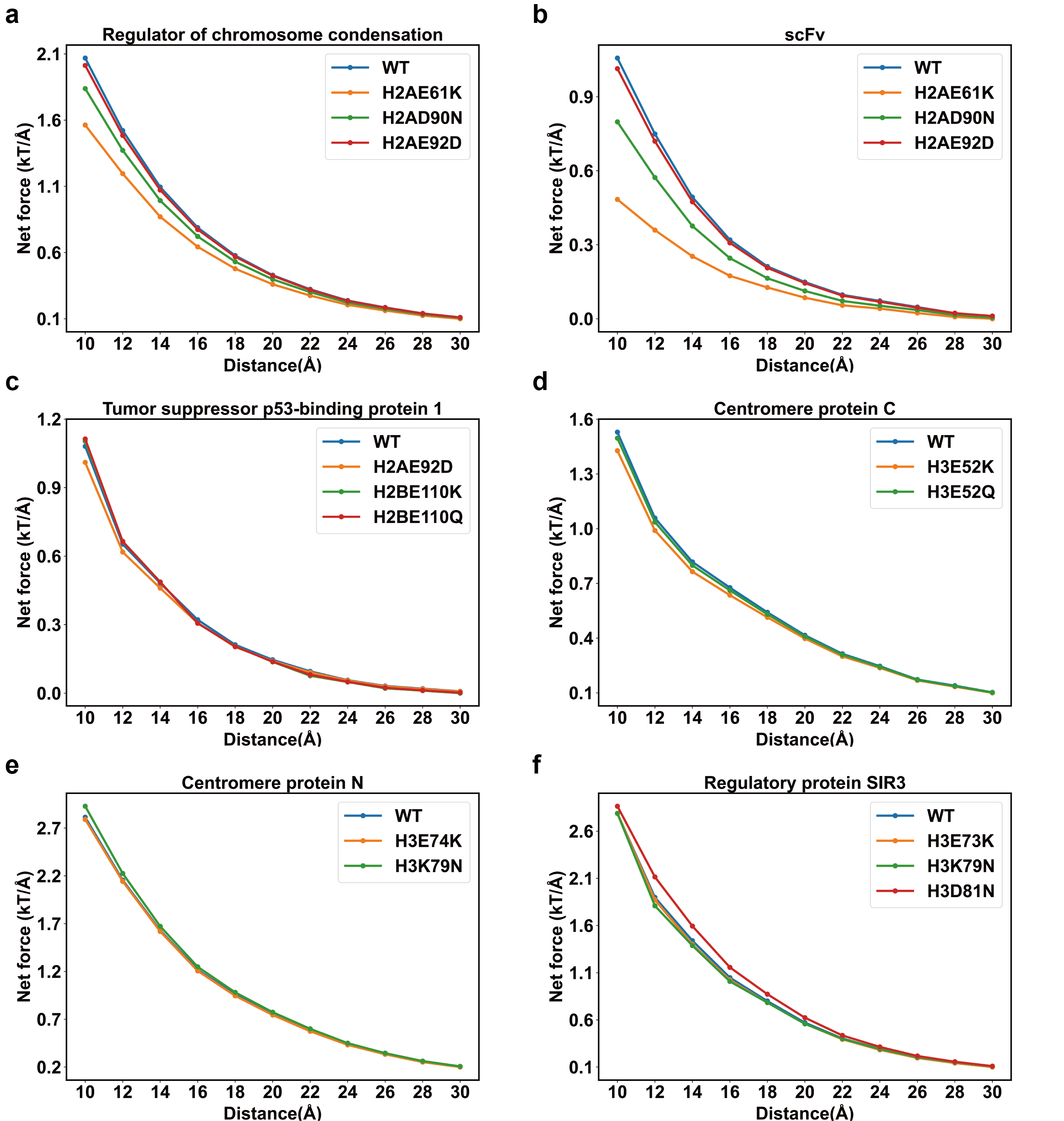

**Supplementary Figure 5.** Comparison of the magnitudes of electrostatic forces between the nucleosome and chromatin factors in wild-type and mutant states. The electrostatic forces were computed at various center-of-mass separating distances ranging from 10 to 30 Å, with 2 Å increments, using PDB structures 3MVD, 6E0P, 5KGF, 6MUP, and 3TU4.

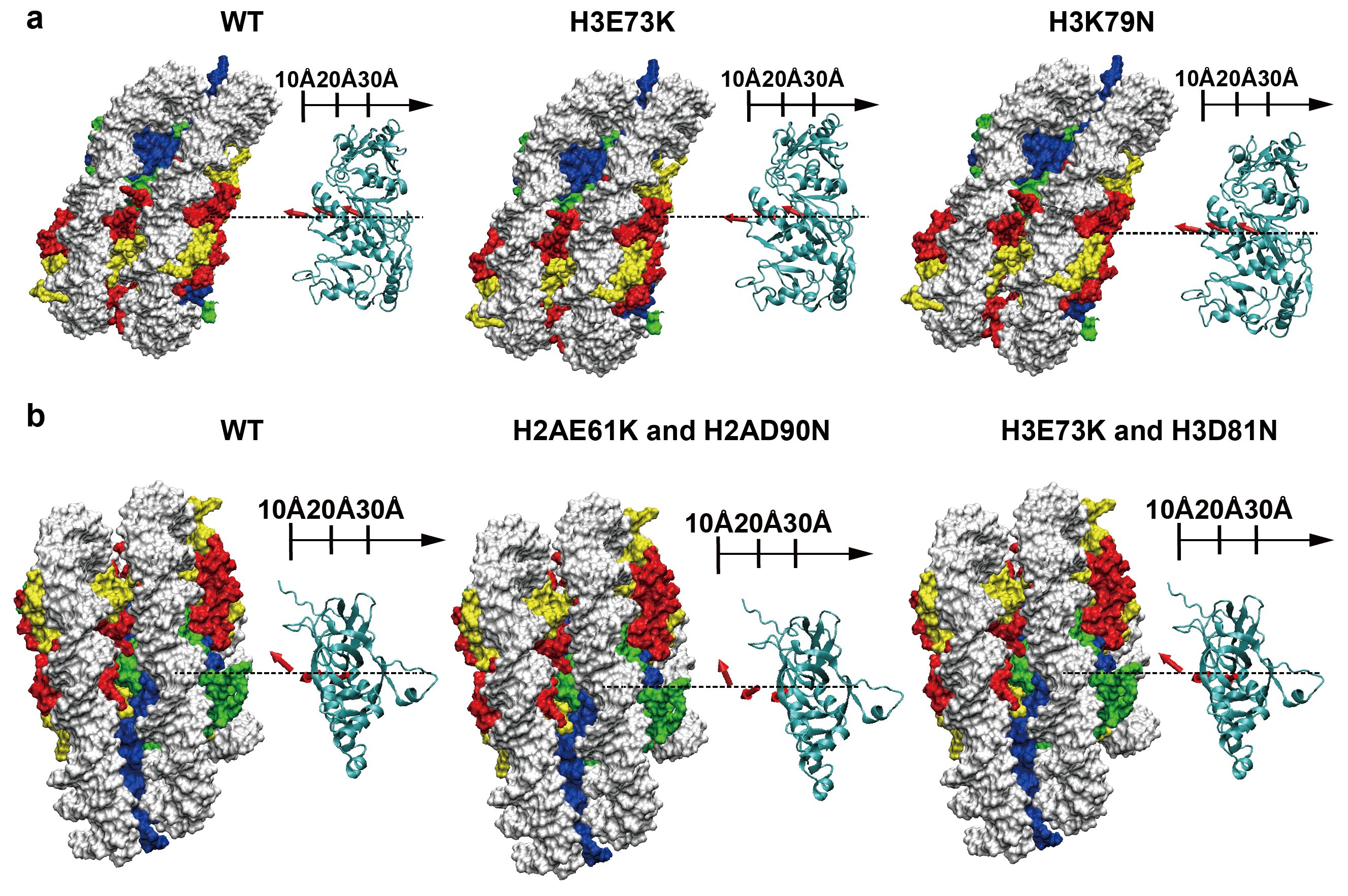

**Supplementary Figure 6.** Effects of histone cancer mutations on nucleosome-chromatin factor long-range electrostatic interactions. (a) and (b) Comparison of the directions of electrostatic forces between nucleosomes and binding proteins (PCR1 above, and regulatory protein SIR3 below) in wild-type and mutant states. The directions of electrostatic interactions are represented by red arrows at separating distances of 10 Å, 20 Å, and 30 Å.

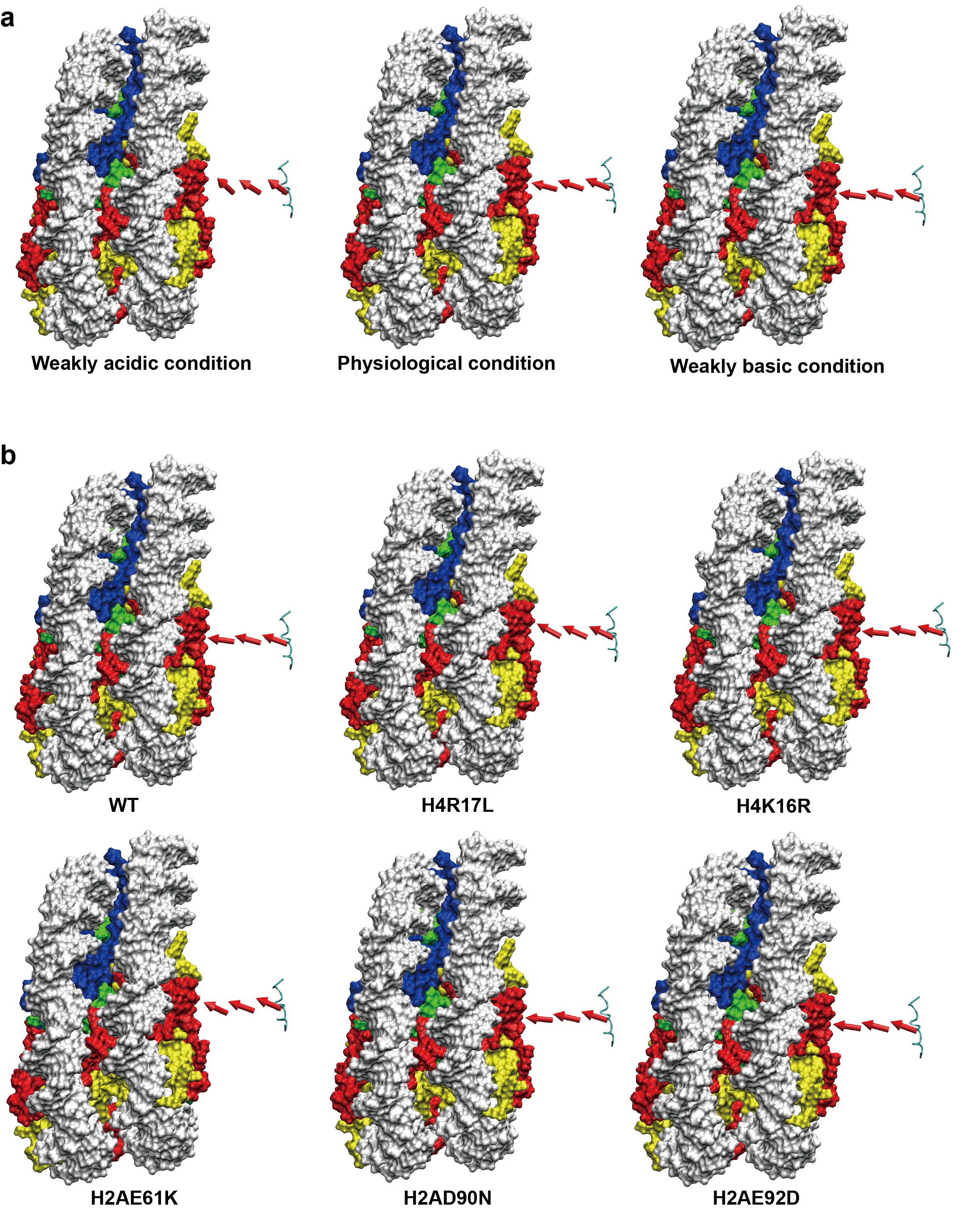

**Supplementary Figure 7.** pH perturbation and histone cancer mutation affect the electrostatic interactions between H4 tail and acidic patch. (a) Comparison of the directions of electrostatic forces between nucleosome and histone H4 tail under weakly acidic (pH 5 to 6.5), physiological (pH 6.5 to 7.5), and weakly basic (pH 7.5 to 9) conditions. (b) Comparison of the directions of electrostatic forces between nucleosomes and H4 tails in wild-type and mutant states. The directions of electrostatic interactions are represented by red arrows at separating distances of 10 Å, 20 Å, and 30 Å.

| PDB ID | 3TU4, 6FTX, 7UV9, 7VVU, 7ZS9, 6T7B, 5KGF, 6G0L, 7Y7I, 8G8G, 7ZSA, 7EG6, 7ENN, 6RYU, 5GTC, 6YOV, 6TDA, 7EGP, 7U50, 6VEN, 6S01, 6T90, 6KW3, 6GEJ, 6LTJ, 6PA7, 7PH6, 7E9F, 6T9L, 7OHA, 8H1T, 7XCT, 6VYP, 7XZZ, 6E0P, 7OH9, 7LYC, 7BWD, 6R1U, 7Y8R, 6NZO, 7XD0, 7LYB, 6Z6P, 6R25, 5MLU, 7D1Z, 6R8Z, 7SCZ, 7W9V, 8GPN, 8GRM, 7EA8, 6R91, 7SSA, 5E5A, 6KIU, 6WKR, 8F86, 6R90, 6QLD, 8AV6, 6PWV, 6PWX, 5X0X, 7TAN, 7YWX, 8ATF, 6KIW, 8DU4, 5X0Y, 6USJ, 6MUP, 6PWF, 7E8D, 3MVD, 6NE3, 6X0N, 7CCQ, 6JYL, 7CRQ, 8H0V, 6T7C |
| --- | --- |

**Supplementary Table 1**. 83 representative nucleosome complex structures for pKa calculations.

| Organisms | PDB |
| --- | --- |
| Homo sapiens | 2CV5,3AFA,5AV6,5X7X,3AV2 |
| Xenopus laevis | 2NZD,3LZ0,3UT9,3UTB,7XFC |
| Drosophila melanogaster | 2PYO,2NQB,6PWE |

**Supplementary Table 2.** 13 high-resolution nucleosome structures for pKa calculations of histone titratable residues.

| PDB ID | Force field and water model | Nucleosome binding protein | Protonation state of histone titratable residues in bound and unbound states | Simulation run | Function of the binding protein |
| --- | --- | --- | --- | --- | --- |
| 3TU4 | AMBER FF14SB for protein and OL15 for DNA，  TIP4PEW water model | Regulatory protein SIR3 | Unbound: H4H18(0), H2BH106(+1) | 3*100ns | Transcriptional silencing |
|  |  |  | Bound: H4H18(+1), H2BH106(0) | 3*100ns |  |
| 5KGF |  | Tumor suppressor p53-binding protein 1 | Unbound: H2BH106(+1) | 3*100ns | PTM readers, DNA repair |
|  |  |  | Bound:H2BH106 (0) | 3*100ns |  |
| 6E0P |  | Single chain antibody fragment | Unbound:  H2AE61(-1), H2AE92(-1) | 3*100ns | Kinetochore components |
|  |  |  | Bound:  H2AE61(0), H2AE92(0) |  |  |
| 6LTJ |  | SWI/SNF-related matrix-associated actin-dependent regulator of chromatin subfamily B member 1 | Unbound:  H2AE64(-1), H2AE72(-1) | 3*100ns | Chromatin remodelers |
|  |  |  | Bound:  H2AE64(0), H2AE72(0) |  |  |
| 6PA7 |  | DNA (cytosine-5)-  methyltransferase 3A/B | Unbound:  H2AE64(-1), H2AE92(-1), H2BH106(+1) | 3*100ns | DNA methyl-transferases |
|  |  |  | Bound:  H2AGLH64(0), H2AGLH92(0), H2BHIS106(0) |  |  |
| 6TDA |  | SWI/SNF chromatin remodeler RSC | Unbound:  H2AE61(-1), H2AE102(-1), H2BH106(+1) | 3*100ns | Chromatin remodelers |
|  |  |  | Bound:  H2AE61(0), H2AE102(0), H2BH106(0) |  |  |
| 7D1Z |  | Isoform 2 of N-lysine methyltransferase KMT5A | Unbound:  H4K20(+1), H4H18(+1) | 3*100ns | PTM writers |
|  |  |  | Bound: H4K20(0), H4H18(0) |  |  |

**Supplementary Table 3.** Summary of simulation runs of seven nucleosome complex structures to analyze the effects of the protonation states of histone titratable residues on nucleosome-partner protein interactions. The assigned net charge of each residue was indicated in different protonation state.

| Histone type | Residue number | Predicted pKa | Standard deviation | Experimental pKa |
| --- | --- | --- | --- | --- |
| H2A | E55 | 5.60 | 0.44 | 4.3 |
|  | E60 | 4.40 | 0.22 | 4.5 |
|  | E63 | 3.59 | 0.28 | 4.5 |
|  | E90 | 4.66 | 0.12 | 4.5 |
|  | E91 | 3.55 | 0.29 | 4.4 |
| H2B | H46 | 6.62 | 0.26 | 5.9 |
|  | E102 | 3.49 | 0.15 | 3.7 |
|  | H106 | 6.40 | 0.04 | 6.5 |
|  | E110 | 3.94 | 0.61 | 4.0 |

**Supplementary Table 4.** Comparison of predicted pKa values of nine histone residues around the acidic patch of the nucleosome with experimentally measured values. The predicted pKa values were calculated from 13 high-resolution nucleosome structures (see Supplementary Table 2). The values were averaged over all structures for each residue.

| Histone type | Residue | Location |
| --- | --- | --- |
| H2A | E61, K95 | Histone-histone binding interface |
| H2B | E71, R72, E76, R92, E93 |  |
| H3 | E59, E73, E97, E105, D106 |  |
| H4 | D68, E74, H75, D85, R95 |  |
| H2A | R17, R29, H31, E41 | Histone-DNA binding interface |
| H2B | R33, E35 |  |
| H3 | R42, E50, D81 |  |
| H4 | R35, R36, R45 |  |

**Supplementary Table 5.** Location of histone titratable residues with significant pKa shifts (|ΔpKa| > 1.5).

| Histone residue | Binding interface | SASA_nuc_ | SASA_dimer_ | SASA_nuc_ -SASA_dime_ |
| --- | --- | --- | --- | --- |
| H3E50 | HDI | 23.98 | 31.79 | -7.81 |
| H3E97 | HHI | 6.54 | 5.77 | 0.78 |
| H3E105 | HHI | 73.00 | 94.48 | -21.48 |
| H3D106 | HHI | 25.29 | 64.27 | -38.99 |
| H4D68 | HHI | 13.29 | 2.01 | 11.28 |
| H4E74 | HHI | 75.09 | 86.13 | -11.04 |
| H4H75 | HHI | 5.19 | 165.83 | -160.64 |
| H2AH30 | HHI | 14.60 | 31.10 | -16.51 |
| H2AE40 | HDI | 130.42 | 152.95 | -22.53 |
| H2AE60 | HHI | 56.76 | 51.46 | 5.30 |
| H2AD89 | HHI | 45.55 | 78.64 | -33.09 |
| H2BH46 | HHI | 11.94 | 16.51 | -4.57 |
| H2BE68 | HHI | 67.33 | 94.97 | -27.63 |
| H2BE73 | HHI | 4.71 | 28.96 | -24.25 |
| H2BE90 | HHI | 0.64 | 8.68 | -8.04 |
| H2BH106 | HHI | 88.58 | 84.17 | 4.40 |

**Supplementary Table 6.** Comparison of solvent-accessible surface area (SASA) of histone residues in nucleosome structure to the values in the form of free dimer. HDI denotes histone-DNA binding interface, and HHI indicates histone-histone binding interface.

| pH condition | Protonation state | Acidic residue | Basic residue |
| --- | --- | --- | --- |
| Physiological (6.5-7.5) | Protonated | pKa > 6.5 | pKa > 7.5 |
|  | Deprotonated | pKa < 6.5 | pKa < 7.5 |
| weakly acidic (5-6.5) | Protonated | pKa > 5 | pKa > 6.5 |
|  | Deprotonated | pKa <5 | pKa < 6.5 |
| weakly basic (7.5-9) | Protonated | pKa > 7.5 | pKa > 9 |
|  | Deprotonated | pKa < 7.5 | pKa < 9 |

**Supplementary Table 7.** Determination of the protonation state of histone titratable residue based on the predicted pKa value.

| Histone residue | Net charge in weakly acidic condition  (5< pH <6.5) | Net charge in physiological condition  (6.5< pH <7.5) | Net charge in weakly basic condition  (7.5< pH <9) | Located in the nucleosome surface |
| --- | --- | --- | --- | --- |
| H3E50 | 0 | -1 | -1 | No |
| H3E97 | 0 | 0 | -1 | No |
| H3E105 | 0 | 0 | -1 | Yes |
| H3D106 | 0 | -1 | -1 | No |
| H4D68 | 0 | -1 | -1 | No |
| H4E74 | 0 | -1 | -1 | Yes |
| H2AE41 | 0 | -1 | -1 | No |
| H2AE61 | 0 | -1 | -1 | Yes |
| H2AD90 | 0 | -1 | -1 | Yes |
| H2BH46 | +1 | 0 | 0 | Yes |
| H2BE71 | 0 | 0 | -1 | No |
| H2BE76 | 0 | -1 | -1 | No |
| H2BE93 | 0 | -1 | -1 | No |
| H2BH106 | +1 | +1 | 0 | Yes |

**Supplementary Table 8.** Protonation state of nucleosome residues under different pH conditions.

| PDB | Binding protein | Function of binding protein |
| --- | --- | --- |
| 3TU4 | Regulatory protein SIR3 | Transcriptional silencing |
| 5KGF | TP53-binding protein 1 (53BP1), Polyubiquitin-B | PTM readers, DNA repair |
| 6E0P | Single chain antibody fragment (scFv) | Kinetochore components |
| 6TDA | SWI/SNF chromatin remodeler RSC (RSC) | Chromatin remodelers |
| 7E9F | Origin recognition complex subunit 1 (Orc1) | DNA replication, Meiosis |
| 7LYB | BRCA1-BARD1-UbcH5c | PTM writers |
| 7XCT | DOT1L | PTM writers |
| 8GRM | PRC1 | PTM writers |

**Supplementary Table 9.** Eight representative nucleosome complex structures for analyses of long-range electrostatic forces across different pH ranges.

| Histone residue | Net charge in weakly acidic condition  (5< pH <6.5) | Net charge in physiological condition  (6.5< pH <7.5) | Net charge in weakly basic condition  (7.5< pH <9) |
| --- | --- | --- | --- |
| H3E74 | 0 | 0 | -1 |
| H3D78 | 0 | -1 | -1 |
| H3D82 | 0 | -1 | -1 |
| H4H20 | 0 | 0 | 0 |
| H4K80 | +1 | +1 | 0 |
| H2AE65 | 0 | 0 | -1 |
| H2AD73 | 0 | -1 | -1 |
| H2AE93 | 0 | 0 | -1 |
| H2BH50 | 0 | 0 | 0 |
| H2BE106 | 0 | -1 | -1 |
| H2AE110 | 0 | 0 | 0 |

**Supplementary Table 10.** Changes in the protonation states of histone titratable residues (Bound state with chromatin factors) under different pH conditions

| PDB ID | Binding free energy of the protonation state (complex) | Standard error | binding free energy of the protonation state (nucleosome) | Standard error |
| --- | --- | --- | --- | --- |
| 7D1Z | -123.73 | 5.12 | -90.61 | 7.60 |
| 3TU4 | -95.67 | 3.86 | -68.63 | 12.16 |
| 5KGF | -89.05 | 7.40 | -87.43 | 6.36 |
| 6LTJ | -54.66 | 2.51 | -60.13 | 6.75 |
| 6TDA | -40.48 | 1.41 | -26.12 | 3.59 |
| 6PA7 | -33.84 | 0.96 | -33.02 | 3.87 |
| 6E0P | -30.62 | 5.30 | -24.50 | 7.80 |

**Supplementary Table 11**．Binding free energy between nucleosome and regulatory proteins in different protonation states. The 'protonation state (complex)' denotes the protonation state in the nucleosome-regulatory protein complex, while the 'protonation state (nucleosome)' refers to the protonation state in the individual nucleosome. Unit for binding free energy are in kcal/mol.

| Mutation | Location | Frequency | Cancer type |
| --- | --- | --- | --- |
| H2AE61K | Acidic patch | 3 | Cervical Cancer, Bladder Cancer |
| H2AD90N | Acidic patch | 3 | Prostate Cancer, Small Cell Lung Cancer |
| H2AE92D | Acidic patch | 3 | Mature B-Cell Neoplasms |
| H3E52K | H3α1L1 elbow | 2 | Mature B-Cell Neoplasms, Gastrointestinal Stromal Tumor |
| H3E52Q | H3α1L1 elbow | 4 | Ovarian Epithelial Tumor, Bladder Cancer, Non-Small Cell Lung Cancer, Breast Cancer |
| H3E73K | H3α1L1 elbow | 10 | Non-Small Cell Lung Cancer, Breast Cancer, Ovarian Cancer, |
| H3K79N | H3α1L1 elbow | 4 | Uterine Endometrioid Carcinoma, Endometrial Cancer, Head and Neck Cancer, Endometrial Cancer |
| H3D81N | H3α1L1 elbow | 9 | Breast Cancer, Non-Small Cell Lung Cancer, Bladder Cancer, Renal Cell Carcinoma, Anal Cancer |

**Supplementary Table 12.** A list of recurrent histone cancer mutations induces significant effects on the nucleosome surface electrostatic potentials.

| PDB | Binding protein | Function of binding protein |
| --- | --- | --- |
| 5KGF | TP53-binding protein 1 (53BP1), Polyubiquitin-B | PTM readers, DNA repair |
| 6MUP | Centromere protein N/C | Kinetochore components |
| 3TU4 | Regulatory protein SIR3 | Transcriptional silencing |
| 8GTM | PRC1 | PTM writers |
| 3MVD | Regulator of chromosome condensation | RanGTP gradient signal |
| 6E0P | Single chain antibody fragment (scFv) | Kinetochore components |

**Supplementary Table 13.** Representative nucleosome complex structures for analyses of histone cancer mutations’ effects on electrostatic interactions.

| Mutation | Location | Frequency | Cancer type |
| --- | --- | --- | --- |
| H2AE61K | Acidic patch | 3 | Cervical Cancer, Bladder Cancer |
| H2AD90N | Acidic patch | 3 | Prostate Cancer, Small Cell Lung Cancer |
| H2AE92D | Acidic patch | 3 | Mature B-Cell Neoplasms |
| H4K16R | H4 tail | 2 | Colorectal Cancer, Hepatobiliary Cancer |
| H4R17L | H4 tail | 2 | Prostate Cancer, Non-Small Cell Lung Cancer |
| H4R19C | H4 tail | 3 | Soft Tissue Sarcoma, Endometrial Cancer, Mature B-Cell Neoplasms |

**Supplementary Table 14.** A list of recurrent histone cancer mutations induces significant effects on H4 tail-acidic patch interactions.
